# Complex petal spot formation in the Beetle Daisy (*Gorteria diffusa*) relies on spot-specific accumulation of malonylated anthocyanin regulated by paralogous GdMYBSG6 transcription factors

**DOI:** 10.1101/2023.02.20.529304

**Authors:** Róisín Fattorini, Farahnoz Khojayori, Gregory Mellers, Edwige Moyroud, Eva Herrero Serrano, Roman T Kellenberger, Rachel Walker, Qi Wang, Lionel Hill, Beverley J Glover

## Abstract

*Gorteria diffusa* has elaborate petal spots that attract male bee-fly pollinators through sexual deception but the genetic basis of *G. diffusa* petal spot development is currently unknown. Here we investigate the regulation of pigmentation during spot formation.

We used UHPLC-MS/MS to determine the anthocyanin composition of spots and background pigmentation in *G. diffusa*. Combining gene expression analysis with protein interaction assays we characterised three R2R3-MYB genes regulating anthocyanin production in *G. diffusa* spots.

We found that cyanidin 3-glucoside pigments *G. diffusa* ray floret petals. Unlike other petal regions, spots contain a high proportion of malonylated anthocyanin. We identified three paralogous subgroup 6 R2R3-MYB transcription factors that activate the production of petal spot pigmentation. The corresponding genes are upregulated in developing spots and induce ectopic anthocyanin production upon heterologous expression in tobacco. EMSAs and dual luciferase assays suggest that these transcription factors regulate genes encoding three anthocyanin synthesis enzymes: anthocyanidin synthase (GdANS), dihydroflavonol reductase (GdDFR) and malonyl transferase (GdMAT1), accounting for the spot-specific production of malonylated pigments.

Here we provide the first molecular characterisation of *G. diffusa* spot development, showing that the elaboration of complex spots begins with accumulation of malonylated pigments at the base of ray floret petals, positively regulated by three subgroup 6 R2R3-MYB transcription factors.

## Introduction

The huge variety of flower colouration present in the natural world is produced by pigmentation and structural effects (Moyroud and Glover 2017). One function of brightly coloured flowers is to attract animal pollinators and several studies have demonstrated that changes in flower colour impact pollinator selection in groups including moths, bees, and hummingbirds (Bradshaw Jr and Schemske 2003; Hoballah et al. 2005; Papiorek et al. 2016; Sheehan et al. 2012). In many species pigmentation patterning occurs within the flower, such as colour differences between floral organs, stripes on petals, and petal spots (Eckhart et al. 2006; Gaskett 2011; Leonard and Papaj 2011; Moeller 2005; Shang et al. 2011). Petal spots are composed of pigmented cells that form discrete aggregations which contrast with the background petal colouration. Spots have evolved independently several times and are present in multiple plant families including Orchidaceae, Fabaceae, Liliaceae, and Asteraceae (Martins et al. 2013). Petal spots can attract pollinators by forming visual or tactile nectar guides (Leonard and Papaj 2011), by increasing the conspicuousness of flowers (de Jager et al. 2017), and by prompting aggregation or mating behaviours (Ellis and Johnson 2010; Johnson and Midgley 1997).

In many species, anthocyanins, which produce red, pink, purple and blue colours (Grotewold 2006; Zhao and Tao 2015), are the major class of petal spot pigments (e.g. Abe et al. 2002; Banba 1967; Martins et al. 2013; Yuan et al. 2014). The anthocyanin biosynthesis pathway has been well characterised in a broad range of species and begins with chalcone synthase catalyzing the formation of tetrahydroxychalcone (Grotewold 2005, 2006; Winkel-Shirley 2001). Downstream enzymes in the anthocyanin synthesis pathway are encoded by late biosynthesis genes (LBGs), namely *DFR* (*DIHYDROFLAVANOL 4-REDUCTASE*), *ANS* (*ANTHOCYANIN SYNTHASE*), and *UFGT* (*UDP-FLAVONOID GLUCOSYL TRANSFERASE*).

R2R3-MYB transcription factors are known regulators of anthocyanin biosynthesis in many systems (Albert et al. 2011; Grotewold 2006; Schwinn et al. 2006). These MYBs contain an activation or repression domain at the C-terminus and two MYB repeats (R2 and R3) at the N-terminus, which together constitute a binding domain for specific DNA sequences (Dubos et al. 2010). Based on these conserved regions R2R3-MYBs have been categorised into subgroups and the regulation of anthocyanin biosynthesis is strongly associated with subgroups 5, 6, and 7 (Feller et al. 2011; Stracke et al. 2001). LBGs are often regulated by subgroup 6 R2R3-MYB transcription factors that form an MBW complex with a basic helix-loop-helix (bHLH) transcription factor and a WD-repeat (WDR) protein (Dubos et al. 2010; Gonzalez et al. 2008; Petroni and Tonelli 2011; Quattrocchio et al. 2006; Ramsay and Glover 2005; Schwinn et al. 2006; Spelt et al. 2000; Stracke et al. 2007).

The development of robust petal patterns requires molecular mechanisms that enable adjacent cells to adopt distinct fates and to restrict pigment production and accumulation to certain regions of the epidermis (recently reviewed in Fairnie et al. 2022). The strict spatial control of anthocyanin synthesis is often attained through transcriptional regulation. Multiple R2R3-MYB genes can regulate floral anthocyanin production within a single species, for example in *Antirrhinum majus* (Schwinn et al. 2006) and *Petunia* (Albert et al. 2011; Albert et al. 2014; Gerats et al. 1985; Gerats et al. 1984). The role of R2R3-MYB transcription factors in regulating petal spot pigmentation has been demonstrated in several species. The tepals of *Lilium* spp. develop spots pigmented by anthocyanin. The *LhMYB12- Lat* allele encodes a protein that regulates ‘splatter’ spot formation, a second *LhMYB12* allele is involved in background pigmentation, while pigmentation in ‘raised’ spots is regulated by *LhMYB6* (Yamagishi et al. 2010). In *Clarkia gracilis, CgMYB1* expression is spatially restricted to the spotted petal region and the corresponding protein activates expression of the gene encoding dihydroflavonol-4-reductase (DFR) leading to spot production, with spot position on the corolla dependent upon which *CgMYB1* allele is present (Martins et al. 2013; Martins et al. 2017). Using experimental data combined with mathematical modelling Ding et al. (2020) demonstrated how floral anthocyanin activators and repressors may interact with each other to form reaction-diffusion systems capable of producing repeated spot patterns, such as those in yellow monkeyflowers (*Erythranthe guttata*, formerly *Mimulus guttatus*).

The sexually deceptive petal spots of *Gorteria diffusa* are highly complex, requiring coordinated regulation of multiple pigment pathways and different cell types (Thomas et al. 2009). Thus, they constitute an excellent system to start unravelling the developmental mechanisms regulating the production of sophisticated motifs on petal surfaces. *Gorteria diffusa* Thunb. is a daisy species (Asteraceae) endemic to southern Namibia and to the South African Richtersveld, Namaqualand and the Western Cape (Duncan and Ellis 2011; Roessler 1959). The Asteraceae capitulum is a compressed inflorescence containing actinomorphic disc florets (flowers) in a central cluster surrounded by zygomorphic ray florets. In *G. diffusa* the long ventral petals of the ray florets are fused into a ligule and dark petal spots, pigmented by anthocyanin, develop across this ligule (Bello et al. 2013; Karis 2007; Thomas et al. 2009). There are geographically defined differences in the floral phenotype of *G. diffusa*, with high levels of capitulum trait variation across a narrow endemic range. Geographically proximal populations have similar phenotypes and sharp boundaries occur where floral traits vary dramatically in adjacent populations. Following quantitative phenotypic investigations, *G. diffusa* populations were assigned to floral morphotypes, defined as geographically discrete floral forms identifiable by differences in capitulum phenotypes. The number of spotted ray florets and the micromorphology of the spot also adds to the phenotypic variation found between morphotypes (Ellis and Johnson 2009). *G. diffusa* petal spot types can be loosely categorised into ‘simple’ dark basal spots present on all ray florets, spots on all ray florets with a raised appearance due to the curvature of ray floret lamina and raised spots that tend to be present on one to four ray florets per capitulum. Raised spots are generally more structurally complex, and in most floral morphotypes they are deeply pigmented three-dimensional elaborations of the petal epidermis comprised of multiple specialised epidermal cell types: green interior cells, white highlight cells, and multicellular black papillae (Thomas et al. 2009). These ‘complex’ *G. diffusa* petal spots are unusual compared to those of other daisies and eudicot species which tend to have less intricate phenotypes (Figure 1) (Thomas et al. 2009).

**Figure 1.**
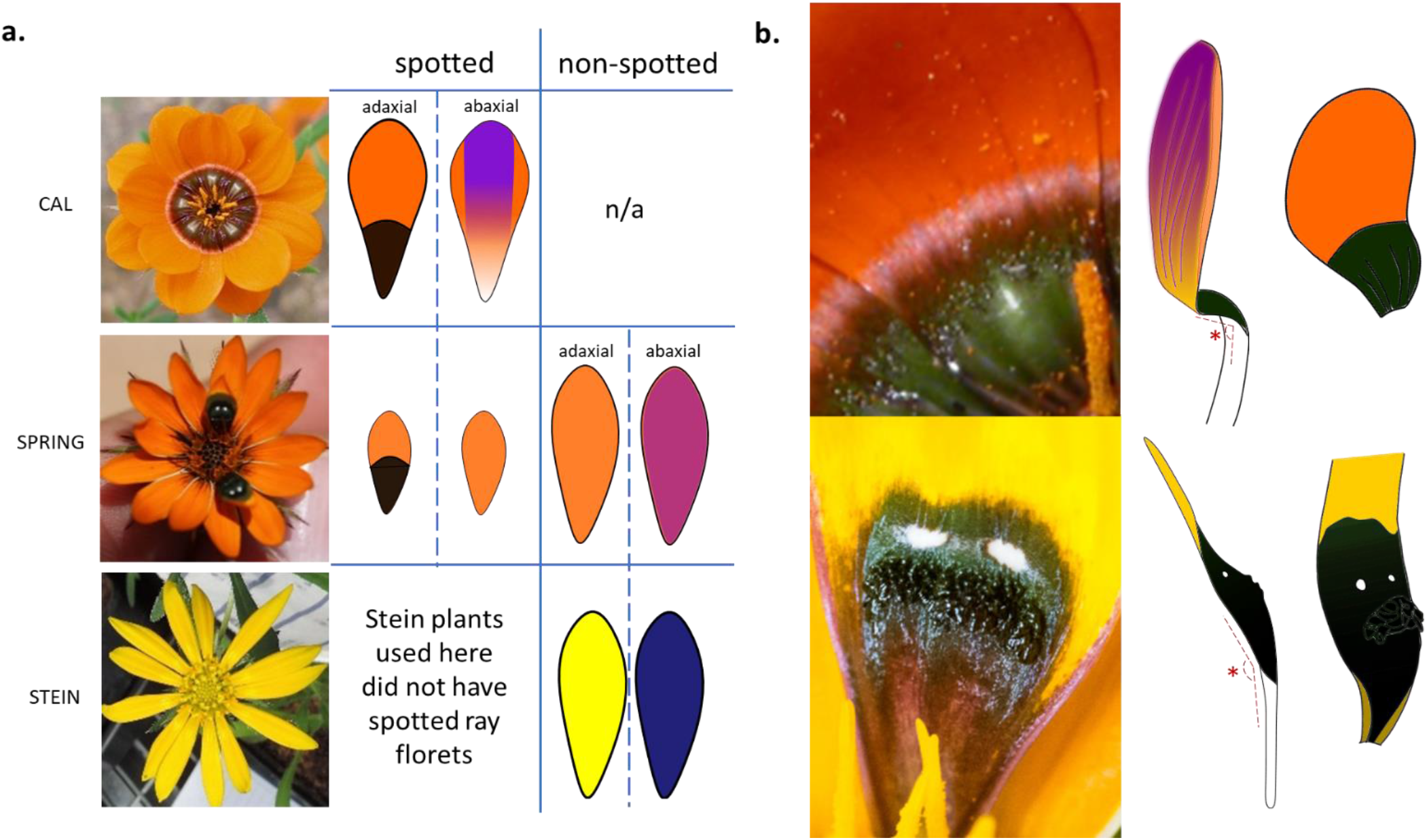
Ray floret, petal, and spot morphology in the three selected morphotypes (Cal, Spring, and Stein) of *G. diffusa*. a) Capitulum phenotypes, ray floret types, and typical petal colouration. Cal capitula are ‘ringed’ with a complex spot forming across the petals of every ray floret. Spring capitula contain 1 - 4 ray florets which develop complex spots – these ray florets have a different overall morphology compared to ‘non-spotted’ Spring ray florets. Some Spring individuals have a small patch of black pigment at the base of all ‘non-spotted’ ray florets (not shown here). Some Stein individuals develop spots on multiple ray florets, the Stein plants used here did not develop any spots on any capitula. b) Complex tridimensional architectures of *G. diffusa* petal spots. Close-up view of Cal (top) and Spring (bottom) petal spots and profile view diagrams of the spotted ray florets. * is the angle of ray floret presentation as defined by Ellis et al. (2014).

*G. diffusa* attracts a bee fly pollinator, *Megapalpus capensis* (Wiedemann), that exhibits different behaviours on the daisy capitula: feeding on pollen and nectar, brief inspection visits, and active attempts to copulate with the spots (Ellis and Johnson 2010; Johnson and Midgley 1997). These pseudocopulatory responses are only observed on specific morphotypes (Spring, Buffels, and Nieuw) that have raised complex spots on a subset of ray florets (Ellis and Johnson 2010; Thomas et al. 2009). The sexually deceptive petal spots are thought to mimic females, as only male flies attempt copulation (Ellis and Johnson 2010; Johnson and Midgley 1997). Sexual deception may increase outcrossing rates as the behaviour of the fly could enable more effective pollen transfer between individual plants (Ellis and Johnson 2010).

In this study we isolated R2R3-MYB transcription factors in *G. diffusa* that promote and spatially restrict the accumulation of anthocyanin to developing petal spots. We first assessed the distribution and composition of anthocyanin pigment within ray floret petals and discovered that malonylated anthocyanins are near exclusive to the spotted region. Multiple *G. diffusa* morphotypes were used to gain robust comparative information on gene expression patterns in spotted and non-spotted ray floret tissue. Protein properties were comparatively investigated using heterologous systems and biochemical assays. Of the four R2R3-MYB transcription factors isolated (*GdMYBSG6-1* to *GdMYBSG6- 4*), three were likely regulators of petal spot anthocyanin production based on expression data. We found that heterologous expression in *Nicotiana tabacum* L. of all three candidates under a strong constitutive promoter was sufficient to activate anthocyanin production. In parallel, we isolated the promoter regions of *ANS*, *DFR*, and *MAT1 (MALONYL TRANSFERASE)* homologues in *G. diffusa*. We identified GdMYBSG6 binding sites possibly acting as cis-regulatory elements in these genomic regions and showed that the *GdDFR* promoter was sufficient to allow GdMYBSG6 to activate transcription of this late biosynthesis gene.

## Materials and Methods

### Plant material

Seeds were collected from wild populations of *G. diffusa* and *G. personata* in the Northern Cape of South Africa under research and collecting permits issued by the Northern Cape Department of Environment and Nature Conservation and Cape Nature. *N. tabacum* and *N. benthamiana* were grown from laboratory lines maintained by selfing. Plants were grown from seed at Cambridge University Plant Growth Facility under 16 hrs light/ 8 hrs dark, 20°C, 60% humidity. Each infructescence of *G. diffusa* was submerged in water overnight and then seeds were planted in two parts Levington’s M3 potting compost and one part horticultural sand (Westland horticulture).

### Anthocyanin pigment extraction and quantification

Samples were collected from *G. diffusa* and *N. tabacum* for anthocyanin extraction. Mature ray floret petals were removed from several *G. diffusa* plants, dissected, and pooled according to petal segment type. Sepals, petals, and anthers were removed from individual *N. tabacum* flowers and the anthocyanin content of each floral organ separately determined.

Samples were snap frozen in liquid nitrogen and pulverised in a tissue lyser at 30 Hz for 30 secs (Qiagen TissueLyser II). Anthocyanin pigments were extracted by adding acidic methanol (1% (v/v) 1M HCL), then samples were vortexed and shaken overnight in the dark. The supernatant was removed and stored at -20°C. The process was repeated, with acidic methanol added to the pelleted material. The anthocyanin levels in the combined supernatants were detected using a spectrophotometer measuring absorbance at the wavelengths A530 and A657. The overall anthocyanin content per gram of tissue was determined as described in Methods S1. Separate samples were used for ultra-high performance liquid chromatography - mass spectrometry (UHPLC-MS/MS) analysis. For each *G. diffusa* tissue type three samples were analysed through UHPLC-MS/MS and one petal sample from transgenic *N. tabacum* was also analysed.

UHPLC-MS/MS analyses were conducted in the Biomolecular Analysis Facility at John Innes Centre, Norwich, UK. A Prominence/ Nexera UHPLC system attached to an ion-trap ToF mass spectrometer (Shimadzu) was used to analyse the samples. A 100 × 2.1 mm 2.6μ Kinetex EVO column (Phenomenex) was used for separation using an acetonitrile versus 1% (v/v) formic acid in water gradient (Methods S1) run at 0.5 ml/min and 40°C. UV/ visible absorbance and positive mode electrospray MS were used to detect compounds, with a diode array detector collecting spectra from 200 - 650nm at 6.25 spectra/sec and a time constant of 0.08 secs. MS spectra were collected from *m/z* 220 – 2000 and MS2 spectra from *m/z* 50 - 2000 with a fixed ion accumulation time of 20 msec, an isolation width of *m/z* 3.0, 50% collision energy and 50% collision gas. The conditions of the spray chamber were as follows: 250°C curved desorbation line, 300°C heat block, drying gas ‘on’, and 1.3 l/min nebuliser gas. Compounds were identified by assessing the masses of the anthocyanidin with no decoration in full MS data, matching the peaks in the photodiode array detection that absorb at approximately 525nm and checking for anthocyanin masses. Having found a peak based on a particular anthocyanin reference compound, the full MS spectrum was used to look for coeluting masses. Mass differences between the fragments using either source fragmentation or MS2 data was used to determine which decorations they likely corresponded to. Total peak areas for each sample (automatically calculated by the software) were cross-referenced with anthocyanin quantification through spectrophotometry and the data were found to be highly consistent regarding proportional differences between tissue types. The proportion of anthocyanin within the sample accounted for by each compound was calculated using peak areas relative to total peak areas. These proportions were then multiplied by the mean total anthocyanin concentration for the identical tissue type determined through spectrophotometry and calculated as described in Methods S1. For subsequent comparisons the data were grouped according to whether or not malonyl residues were present.

### Isolation of candidate genes and promoter regions

*GdMYBSG6-1* and *GdMYBSG6-2*, along with partial sequences of *GdDFR* and *GdANS*, were identified from previously published transcriptomic data of *G. diffusa* spotted ray florets (Walker 2012). *GdMYBSG6-3* was amplified using primers designed from the *GdMYBSG6-2* sequence and *G. personata MYBSG6* orthologues were isolated using primers designed from *GdMYBSG6* sequences, except for *GpMYBSG6-4* which was found using degenerate primers (Table S1). *GdMYBSG6-4* and *GdMAT1* were subsequently identified from a second transcriptome analysis (Kellenberger unpublished). The *G. diffusa* DNA used for gene hunting was obtained using a CTAB DNA/ RNA extraction procedure on plant tissue that had been snap frozen in liquid nitrogen and ground in a tissue lyser (Qiagen TissueLyser II). RNA was DNase treated using TURBO DNA-*free* (Ambion) according to the manufacturer’s instructions and first strand cDNA synthesis was performed with oligo dT primers using BioScript reverse transcriptase (Bioline). *GdMYBSG6* sequences have been deposited in GenBank under the following accession numbers: OQ434658, OQ434659, OQ434660, OQ434661, OQ434662, OQ434663, OQ434664, OQ434665, OQ434666, OQ434667, OQ434668.

Genome walking was conducted using the Universal GenomeWalker 2.0 kit (Takara Bio) according to the manufacturer’s instructions to isolate *GdMYBSG6-1* and *GdMYBSG6-2* 5’UTRs and the promoter regions of *GdANS* and *GdMAT1*. The full-length of *GdDFR* and promoter regions were isolated using a combination of genome walking and PCR with degenerate primers (Table S1). 3’RACE (Ren et al. 2005) was used to amplify the 3’ ends and 3’UTR of each gene. PCRs used to genome walk and to obtain full length gene sequences indicated that multiple *GdDFR* copies are likely present. Based on these divergent sequences, multiple primer combinations were used to try to amplify the different *GdDFR* variants (Table S1). The full-length coding sequences of *GdMYBSG6-3* and *GdMYBSG6-4* were isolated using primers designed against the UTRs of the other *GdMYBSG6* genes. The genomic coding sequence and upstream region of *GdMAT1* were amplified using primers designed to be specific to *GdMAT1* (Table S1). The availability of raw *G. diffusa* genome sequencing reads (Kellenberger, unpublished) enabled forward primers to be designed to amplify further upstream of *GdANS* and *GdMAT1* coding sequences (Table S1). A single PCR reaction was used to amplify the full length of the promoter region and the corresponding coding sequence, the PCR product was then sequenced using Sanger sequencing (Biochemistry DNA Sequencing Facility, University of Cambridge, UK). Sequencing data were formatted and analysed in Geneious Prime and Benchling (Biology Software). All PCRs were conducted with the proof-reading enzyme Phusion (New England Biolabs) according to the manufacturer’s instructions.

### Phylogenetic analyses

A gene tree was constructed using the coding sequences of Asteraceae subgroup 6 MYBs. Asteraceae sequences were taken from Genbank using BLAST analysis, the lettuce (*Lactuca sativa*) genome (Lettuce Genome Resource, https://lgr.genomecenter.ucdavis.edu/, Reyes-Chin-Wo et al. 2017), the sunflower (*Helianthus annuus* L.) genome (Badouin et al. 2017, https://www.sunflowergenome.org/), and from a literature search for relevant papers (Yue et al. 2018). *GdMYBSG9-1* encodes a subgroup 9 R2R3-MYB (Thomas 2009) and was included as an outgroup. *G. personata* seeds were not available and so only gDNA could be obtained from dried leaf field samples for this species. As such, introns were predicted and removed from *G. personata* sequences by aligning the gDNA sequences with the corresponding *G. diffusa* coding sequences. The cDNA sequences were aligned with transAlign (v1.2) (Bininda-Emonds 2005) using an amino acid alignment to facilitate the cDNA alignment. Default parameters were used with one exception: no gaps were removed, instead of the default of stripping only those flanking the sequence. The model of molecular evolution was selected using PartitionFinder (v2.1.1) (Lanfear et al. 2016) testing all nucleotide models available assuming each of the three codon positions is a data block. The phylogenetic tree was then inferred using maximum-likelihood optimality criteria with RAxML-NG(v.1.1.0) (Kozlov et al. 2019) while calculating bootstrap branch support. The phylogenetic tree was visualised using FigTree (v1.4.4).

### Gene expression analyses

*G. diffusa* ray floret tissue was sampled from three independent biological replicates, each comprising multiple plants, at two developmental stages during spot development (Figure S5). These stages were selected based on the developmental characterisation of the Nieuw morphotype (Thomas et al. 2009). *N. tabacum* petal tissue was harvested at developmental stage one (as defined in Pak Dek et al. 2017) (Figure S9) from each *N. tabacum* transgenic line, this sample consisted of pooled tissue from multiple flowers of one T_1_ individual. *N. tabacum* qRT-PCR analyses were conducted on wild type plants and those transformed with either *35S::GdMYBSG6-1, 35S::GdMYBSG6-2*, or *35S::GdMYBSG6-3* constructs.

RNA used to determine gene expression levels was extracted using a 2% CTAB-based buffer followed by precipitation with 4M lithium chloride method. A TURBO DNA-*free* kit (Ambion) was used to DNase treat the extracted RNA. cDNA was synthesised using SuperScript II reverse transcriptase (Invitrogen) according to the manufacturer’s instructions. qRT-PCR was completed using the Luna Universal qPCR Master Mix (New England Biolabs). The *Elongation Factor 2* homologue (*GdEF-2*) was selected as a suitable reference gene in *G. diffusa* out of four possible candidates through assessment of gene stability across several developmental stages and biological replicates (Table S1) (Chen et al. 2011). Primer pairs for qRT-PCR were designed to be specific to *GdMAT1*, to *GdANS,* or to bind to all variants of *GdDFR* (Table S1). *GdMYBSG6* primers were specific to each gene (Table S1). The reference genes for *N. tabacum*, *Elongation Factor 1* (*NtEF-1*) and *Ubiquitin C* (*NtUBC*), were selected from the literature (Table S1) (Divya et al. 2019; Schmidt and Delaney 2010). *NtDFR* and *NtANS* primer sequences were modified from those used in Yamagishi et al. (2014) (Table S1) and blasted against the *N. tabacum* genome to ensure only *DFR* or *ANS* genes would be amplified. Pairs of primers were selected for qRT-PCR only if efficiency was within the range 1.9 - 2.1. Reactions were conducted in CFX Real-Time PCR machines (CFX384 and CFX Connect) using the following cycle conditions: 95°C for 1 min; 35 cycles of 95°C for 15 secs, 60°C for 30 secs, a plate fluorescence reading; a post-PCR melt curve analysis 60 - 95°C with readings taken at increments of 0.5°C. Data were checked and analysed with the Opticon Monitor software package (BioRad Laboratories, Inc). Expression levels were calculated relative to the reference gene/s using the modified common base method (Ganger et al. 2017) using primer efficiencies calculated for each primer pair.

### Stable transformation of *Nicotiana tabacum*

A pGREEN II plasmid (Hellens et al. 2000) containing a double CaMV 35S promoter sequence was used to constitutively express the *GdMYBSG6* genes in *N. tabacum*. The *GdMYBSG6* coding sequences were amplified from Spring morphotype cDNA using Phusion High-Fidelity DNA Polymerase (New England Biolabs) and cloned into pBLUESCRIPT KS(-) vectors using the *Eco*RV site. These *GdMYBSG6* plasmids and pGREEN II plasmid were digested with restriction enzymes (*Pst*I and *Sal*I) and each *GdMYBSG6* sequence was ligated into a separate pGreen II plasmid (Methods S2). The pGREEN-*GdMYBSG6* plasmid contained *GdMYBSG6-1*, *GdMYBSG6-2* or *GdMYBSG6-3* under the control of a constitutive double 35S CaMV promoter and a 35S terminator sequence. These constructs were transformed into *Agrobacterium tumefaciens* strain GV3101 through electroporation (Gene Pulser XCell by BIO-RAD). Transformed cells were plated onto LB media containing 50mg/l kanamycin and 25mg/l gentamycin and selected following 48 hrs growth at 30°C in the dark. *N. tabacum* cv. Samsun was then stably transformed with these *A. tumefaciens* strains following a modified version of the method of Horsch et al. (1985) (Methods S3), with several independent insertion lines produced for each gene (*GdMYBSG6-1* n = 4, *GdMYBSG6-2* n = 8, *GdMYBSG6-3* n = 8).

Regenerated plants were genotyped via PCR to test for the presence of the transgene. cDNA derived from leaf tissue RNAs and PCRBIO Taq DNA polymerase (PCR Biosystems) were used to check for transgene expression (Table S1, Figure S8). qRT-PCR experiments and anthocyanin quantification (previously described) were conducted on the T_1_ generation of transformants, with wild type controls.

### Production and purification of recombinant GdMYBSG6 proteins

Gibson assembly (Gibson et al. 2009) was used to produce a pETM-11 vector containing a sequence encoding a GdMYBSG6-2 protein tagged by six histidines at each end (Methods S4). Sanger sequencing was used to confirm that the vector sequence was correct. These plasmids were transformed into *E. coli* (rosetta II strain), grown on LB plates containing 30mg/l chloramphenicol and 50mg/l kanamycin. Successful transformation was confirmed through colony PCR and a single colony was used to inoculate 250ml of LB media (containing 30mg/l chloramphenicol and 50mg/l kanamycin). These cultures were grown at 37°C with 180 rpm shaking. Once the OD_600_ reached 0.6, protein production was induced with the addition of 250 µl 1M IPTG, and 500 µl 1M betaine was added. Cultures were then incubated at 30°C with 180 rpm shaking for 2.5 hrs for induction. Following incubation, cultures were centrifuged and the bacteria pellets were frozen at -80°C.

Protein pellets were resuspended in lysis buffer (Methods S5) and sonicated on ice at 20 amps for a duration of 30 secs five times. After centrifugation, the supernatant was loaded onto a column containing 2 ml of Co-NTA resin (WorkBeads^TM^ 40 Co-NTA, Bio-Works) pre-balanced in loading buffer (Methods S5) and the flow through was collected. For the wash step, 20 ml of loading buffer was run through the column followed by 20 ml of filtration buffer 1 (B1) (Methods S5). Finally, filtration buffer 2 (B2) (Methods S5) was added incrementally, and 1ml elution fractions were collected. An aliquot of each fraction, along with the non-induced control culture, was mixed with 4x NuPAGE LDS sample buffer, heated at 85°C for 10 mins to denature the protein, and 15 µl were run on a NuPAGE 12% Bis-Tris gel (Invitrogen) alongside 5 µl ladder (PageRuler Plus Prestained protein ladder, 10 – 250kDa). The gel was run in 1x MOPs SDS buffer, stained with Coomassie blue for 30 mins, transferred to destaining solution (Methods S5), and imaged. The fraction containing high concentrations of purified protein was selected (B1), the purified protein was concentrated, and buffer exchanged into storage buffer (Methods S5) using protein concentrators (Perice Protein Concentrator PES, 10K MWCO) according to the manufacturer’s instructions. The solution was then aliquoted, snap frozen in liquid N2, and stored at -80°C.

### Electromobility shift assay

Possible MYB binding motifs were identified in *GdANS*, *GdDFR*, and *GdMAT1* upstream sequences, first searching for DNA motifs found to bind *A. thaliana* subgroup 6 R2R3-MYB transcription factors in yeast one-hybrid experiments (Kelemen et al. 2015) and the AtMYB113 (a subgroup 6 R2R3-MYB) binding motifs identified by O’Malley et al. (2016). Potential binding motifs were also identified using the position weight matrix describing MYB113 binding preferences to DNA (MA1181.1) available from JASPAR (https://jaspar.genereg.net/matrix/MA1181.1/). An additional possible binding site was identified in the *GdANS* and *GdDFR* promoters based on a match with a motif recognised by an *Ipomoea purpurea* MYB (Wang et al. 2013). For each potential motif, an oligonucleotide spanning the 29 bp promoter region with the predicted motif in the centre was designed and a G nucleotide added to the 5’ end. This oligonucleotide was ordered along with a 29 bp reverse complement (excluding the added G) from IDT (Integrated DNA Technologies). The complementary oligos were annealed in a 50 µl reaction (20 µl 40 µM oligo 1, 20 µl 40 µM oligo 1, 5 µl 10x annealing buffer (Methods S5), 5 µl ddH_2_O) heated to 96°C for 6 mins and gradually cooled to room temperature. The annealed oligos were fluorescently labelled by mixing 5 µl 0.8 µM annealed oligos, 2 µl 10x Klenow buffer, 1 µl 8 µM Cy3-dCTP, 1 µl Klenow enzyme (ThermoFisher Scientific), 11 µl H_2_O and incubating at 37°C for 2 hrs followed by enzyme inactivation 65°C for 10 mins.

EMSA gels were made using 1.8 ml acrylamide (29:1), 600 µl 10x TBE, 120 µl 10x (v/v) APS, 12 µl TEMED and 9.6 ml ddH_2_O. Purified protein was thawed on ice and diluted 1:5 with binding buffer (Methods S5). For the binding reaction 8 – 17 µl of diluted protein, 2 µl 100 ng/µl fish sperm DNA (Sigma-Aldrich), 1 µl Cy3-dCTP labelled oligos, and enough binding buffer to make a 20 µl reaction were mixed and incubated on ice for 30 mins, alongside control reactions containing no protein. Gels were pre-run for 30 mins, then samples were added and run at 90V for 60 – 75 mins at 4 °C. Gel imaging was completed on a ChemiDoc MP (Bio-Rad).

### Dual luciferase assays

The pGREEN plasmid containing *GdMYBSG6-2* under the control of a constitutive double 35S CaMV promoter and a 35S terminator sequence, also used in stable transformation of *N. tabacum* (Methods S2), was used as an effector plasmid in a dual-luciferase assay. The reporter construct was obtained by cloning the available *GdDFRp* promoter region (378 bp) into the pGreen II 0800-LUC vector (Methods S6), following double digestion with KpnI and NcoI and subsequent ligation. The promoter was inserted upstream of the firefly derived luciferase reporter gene (Hellens et al. 2005). The pGreen II 0800-LUC vector also contains a *Renilla* derived luciferase reporter under the control of the CaMV 35S promoter, which acts as an internal control to normalize the values of the experimental reporter gene for variation caused by transfection efficiency.

The effector and reporter plasmid were introduced into the *Agrobacterium tumefaciens* strain GV3101. Liquid cultures of LB media containing 50mg/l kanamycin, 50mg/l rifampicin, and 25mg/l gentamycin were inoculated with a transformed *A. tumefaciens* colony. A 3ml LB culture was grown overnight and 1ml of this culture was then added to 9ml of LB which was grown overnight. Cultures were grown at 28°C, shaking at 130 rpm. Cells were then harvested by centrifugation and resuspended in induction media (10 mM MgCl_2_, 10 mM MES pH 5.6, 200 µM acetosyringone) to a final optical density of A_600_ = 0.25. Induced cultures were infiltrated into the abaxial surface of young leaves of four-week-old *N. benthamiana*. Four infiltrations per leaf were completed, with media containing a single reporter plasmid with the *GdDFR* promoter region, plus an effector plasmid encoding GdMYBSG6-2 or an empty pGREEN as a control. For each combination, an injected leaf on a separate plant was treated as a biological replicate. Infiltrated leaves were harvested 48 hrs after infiltration and luciferase activity was measured immediately using the Dual-Glo Luciferase Assay kit (Promega). Leaf disks 4 mm in diameter were cut and inserted into white 96 well plates with 75 µl of 1x PBS and 75 µl of luciferase assay reagent. Firefly luminescence was measured on the ClariostarPlus (BMG Lab Technologies) after 15 mins. Afterwards, 75 µl of Stop & Glo reagent was added to the samples, samples were incubated for 15 mins, and the *Renilla* luminescence was measured as an internal control. Results were expressed as the ratio of firefly to *Renilla* luciferase activity.

### Data analysis

Calculations were completed in excel and graphical representations of the results were completed in R (packages:ggplot2 (Wickham 2016)). Statistical modelling was completed using R packages multcomp (Hothorn et al. 2008), ggfortify (Horikoshi & Tang 2018; Tang et al. 2016), nlme (Pinheiro et al. 2013), and lsmeans (Lenth 2016). Statistical analyses used were as follows: Linear mixed effects models (with plant ID added as a random effect) and Tukey’s tests for comparing *G. diffusa* anthocyanin concentrations; ANOVA and Tukey’s tests/ FDR method for comparing *G. diffusa* qRT-PCR data, anthocyanin concentration and gene expression between *N. tabacum* GdMYBSG6 transformants; a Pearson correlation test was used to compare *N. tabacum* transgene expression levels with those of *NtANS* and *NtDFR*; a linear mixed model (transgene expression level as random effect) and FDR method were used for comparing *NtANS* expression levels in transformed *N. tabacum*. Data were transformed were necessary to ensure that assumptions of the statistical models were met. UHPLC-MS/MS data analyses were performed as described in Method S1.

## Results

### Morphological characterisation of ray florets in *G. diffusa* morphotypes differing in spot occurrence

To sample across the range of natural variation, individuals from *G. diffusa* morphotypes Calendula (Cal), Springbok (Spring), and Steinkopf (Stein) were used in this study (Figure 1a). These morphotypes are known to be diploid from karyotyping (Thomas 2009) and, interestingly, Cal and Spring capitula induce differential behavioural responses in *M. capensis* – the major pollinating species (Ellis and Johnson 2010). Spots form across the fused petals of a ray floret in all three morphotypes but the frequency of spotted florets varies between each morphotype. In Cal each ray floret produces a spot creating a ringed capitulum phenotype. In Spring one to four non-adjacent ray florets develop spots varying between capitula within an individual. Spring ray florets lacking spots are larger and can have different background colouration than spotted counterparts (Figure 1a, Figure S1). Occasionally, within some Spring individuals, the non-spotted ray florets all display a basal patch of dark pigment – here referred to as a ‘mark’ (Figure S1). Petal spots can sometimes develop on a subset of ray florets in some Stein individuals, but Stein plants used here did not have any spotted capitula.

Cal and Spring petal spots are vibrantly coloured and textured creating a complex and shimmering spot phenotype. In Cal the overall spot colouration is deep green, while in Spring spots appear black/ purple/ green (Figure 1b). The shape of the spotted ray florets creates a raised presentation contributing toward the 3D elaboration of the petal spot. The ‘angle of ray floret presentation’ (Figure 1b) is a greater contributor to the raised appearance of Cal spotted ray florets compared to those of the Spring morphotype (Ellis et al. 2014; Mellers 2016). All three focal morphotypes have purple or black pigmentation on the abaxial side of the ray floret (Figure S2). The extent of this pigmentation varies between individual plants grown in controlled glasshouse environments and between individuals from the same wild populations. Cal and Stein tend to have consistently darker pigmentation extending over more of the abaxial petal area than Spring non-spotted ray florets, while the extent of Spring abaxial pigmentation appears more variable. Dark pigmentation, characteristic of anthocyanins, was largely absent from the abaxial side of both spotted ray floret petals in Spring and the spotted portion of Cal ray florets – this was highly consistent between capitula and individuals (Figure S2). As such, only the ray florets/ ray floret regions that are exposed when flowers are closed have abaxial anthocyanin pigmentation, possibly reducing flower conspicuousness as an anti-herbivore mechanism (Kemp and Ellis 2019). In summary, colouration consistent with anthocyanin pigmentation occurs within the petal spots of Spring and Cal and on the abaxial side of non-spotted petal tissue of both morphotypes.

### Cyanidin accounts for ray floret petal pigmentation and malonylated cyanidins are found predominantly in petal spots

To complement the qualitative characterisation of petal spot and ray floret background pigmentation, the concentrations of anthocyanins in spotted and non-spotted ray floret petal regions (Figure 2a) were quantified through spectrophotometry. Spring spots had a significantly higher anthocyanin content than any other tissue type (Cal non-spotted: z = 3.38, p = 0.009; Stein non-spotted: z = 4.58, p < 0.001; Cal spot: z = 4.66, p < 0.001; Spring mark: z = 7.02, p < 0.001; Spring non-spotted: z = 9.00, p < 0.001) (Figure 2b, Table S2). Cal spots had equivalent anthocyanin concentrations to Cal and Stein non-spotted petal tissue, which have dark abaxial pigmentation. The non-spotted part of Cal ray floret petals contained significantly more anthocyanin than Spring non-spotted ray floret petals (z = 3.57, p = 0.005), potentially due to greater abaxial pigmentation in Cal, and anthocyanins could also contribute to the adaxial deep orange colouration of this morphotype.

**Figure 2.**
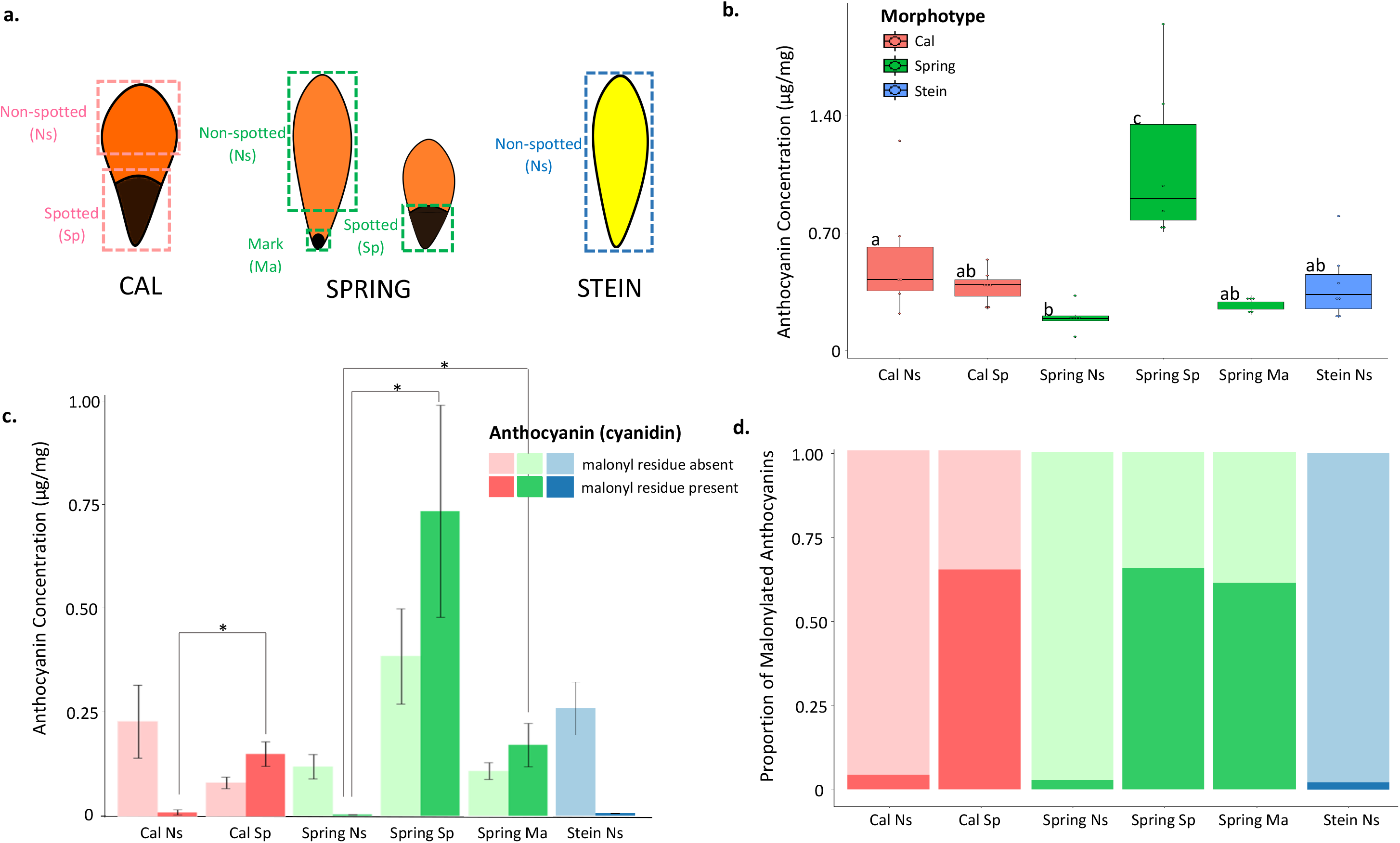
Anthocyanin content of *G. diffusa* ray florets in the morphotypes Cal, Spring, and (non-spotted) Stein. a) Schematics of typical ray florets from each morphotype indicating regions used for pigment extraction (dashed boxes). ‘Spring Mark’ is a small patch of pigment located at the base of non-spotted ray florets in some Spring individuals. b) Anthocyanin content (μg cyanindin-3-glucoside equivalent per mg of fresh tissue) for tissue types depicted in a. Median values and 25/75% quantile +/- 1.5 * interquartile range are indicated. Individual data points are represented by black dots. Sample size n = 5 - 7, where n represents pooled tissue from a single individual. Boxes that do not share letters are significantly different from one another (according to linear mixed effects model and Tukey’s test). c) Summary HPLC-MS analysis results. Approximate anthocyanin content for each tissue type is shown, grouped according to whether a malonate residue is present or absent. Sample size n = 3, where n represents pooled tissue from a single individual, error bars are +/- s. d. * indicates p < 0.0001 d) Proportion of anthocyanin for each sample type that contains a malonate residue colour coded according to the key in c).

UHPLC-MS/MS analyses were conducted to identify the anthocyanins present in the different *G. diffusa* ray floret petal regions from the three morphotypes. A major peak was detected at 5.059 mins in the chromatograms showing UV absorbance within every sample. For the non-spotted regions this was the only major peak (Table S2, Figure S3). The mass of the compound was 449 and collision-induced dissociation (CID) of the [M]_+_ ion at m/z 449 formed a base peak at m/z 287, corresponding to aglycone cyanidin and a mass loss of 162 suggests loss of a glucose moiety (Figure S4, Table S3). Cyanidin 3-glucoside was used as a reference compound by Schütz et al. (2006) during UHPLC-MS/MS analyses on acidic methanol extractions from *Cynara scolymus* L. (Asteraceae). The MS-MS analysis results of their study were identical to the major peak here identified and, as such, we tentatively identify our major peak as cyanidin 3-glucoside.

Two additional major peaks were present in the UV absorbance chromatograms of all petal samples from spotted or marked regions (Cal spot, Spring spot, Spring mark). The retention times of these peaks were 7.133 mins and 8.283 mins, and both had m/z 287 and m/z 449 consistent with a cyanidin glucoside (Figure S4, Table S3). Compound 4 (Table S3) had a mass loss of 44 (mass of 535 and a peak at m/z 491) corresponding to a likely loss of carbon dioxide from the terminal carboxylic acid group of a malonate. Compound 5 (Table S3) had a mass of 549 and a peak at m/z 517, the mass loss of 100 is appropriate for a methylmalonate. In a subset of samples additional minor peaks were detected in the UV absorbance chromatogram (Table S3). All peaks identified were cyanidin (although characterisation was inconclusive in one case) and contained a glucose moiety, except for one cyanidin that had a pentose sugar moiety.

Cyanidins containing a malonyl group were present at concentrations ≥ 12-fold higher in spotted petal tissue compared to non-spotted petal tissue across morphotypes (Figure 2c). The proportion of anthocyanins within spotted petal tissue that contained a malonyl residue was consistent between the Spring and Cal morphotypes (65%) and between the complex (65%) and simple spot (61%) (‘mark’: formed of anthocyanin pigment with no cellular elaborations on non-spotted ray florets) of Spring (Figure 2d). In non-spotted petal tissue cyanidin containing malonyl residues accounted for 6% or less of the total anthocyanin content. The overall anthocyanin composition of Spring marks was similar to that of Cal and Spring complex spots (Table S2). Cyanidin glucosides with caffeate residues accounted for approximately 4% of the anthocyanins within Spring spots, 2% within Spring marks, and 1% within Stein non-spotted ray florets. However, anthocyanin containing caffeate were completely absent or detectable only in trace amounts within all Cal samples (Table S2).

### *G. diffusa* R2R3-MYB transcription factors (GdMYBSG6) are candidate regulators of petal spot anthocyanin

Four R2R3-MYB transcription factors were identified as candidates for regulating petal spot pigmentation in the Spring morphotype. *GdMYBSG6-1* and *GdMYBSG6-2* were first isolated and found to be upregulated in spotted petal tissue within a *G. diffusa* transcriptome (Walker 2012), implying that they encode possible activators of anthocyanin production. Subsequently, *GdMYBSG6-3* was identified as an additional paralogue through gene hunting via PCR. A second transcriptome (Kellenberger unpublished) recovered a fourth paralogue, *GdMYBSG6-4*, which we decided to investigate due to its sequence similarity to the other three genes. All four genes encode the R2 R3 domains and amino acid motif typical of subgroup 6 R2R3-MYBs, suggesting they are likely representatives of this group (Figure 3a). GdMYBSG6-1, GdMYBSG6-2, and GdMYBSG6-3 have 84 - 91% amino acid conservation. GdMYBSG6-4 is the most divergent paralogue sharing 68 - 69% of its amino acid sequence with the other GdMYBSG6 genes (Figure 3a). The *GdMYBSG6* genes were highly structurally conserved in the Cal and Stein morphotypes and very similar in length to the Spring morphotype counterparts: between morphotypes intron 2 and exon 3 for all four genes had slightly varying lengths, and the length of *GdMYBSG6-4* intron 1 also differed. Only a small section of *GdMYBSG6-2* was isolated in Stein, so this gene was excluded from this comparison.

**Figure 3.**
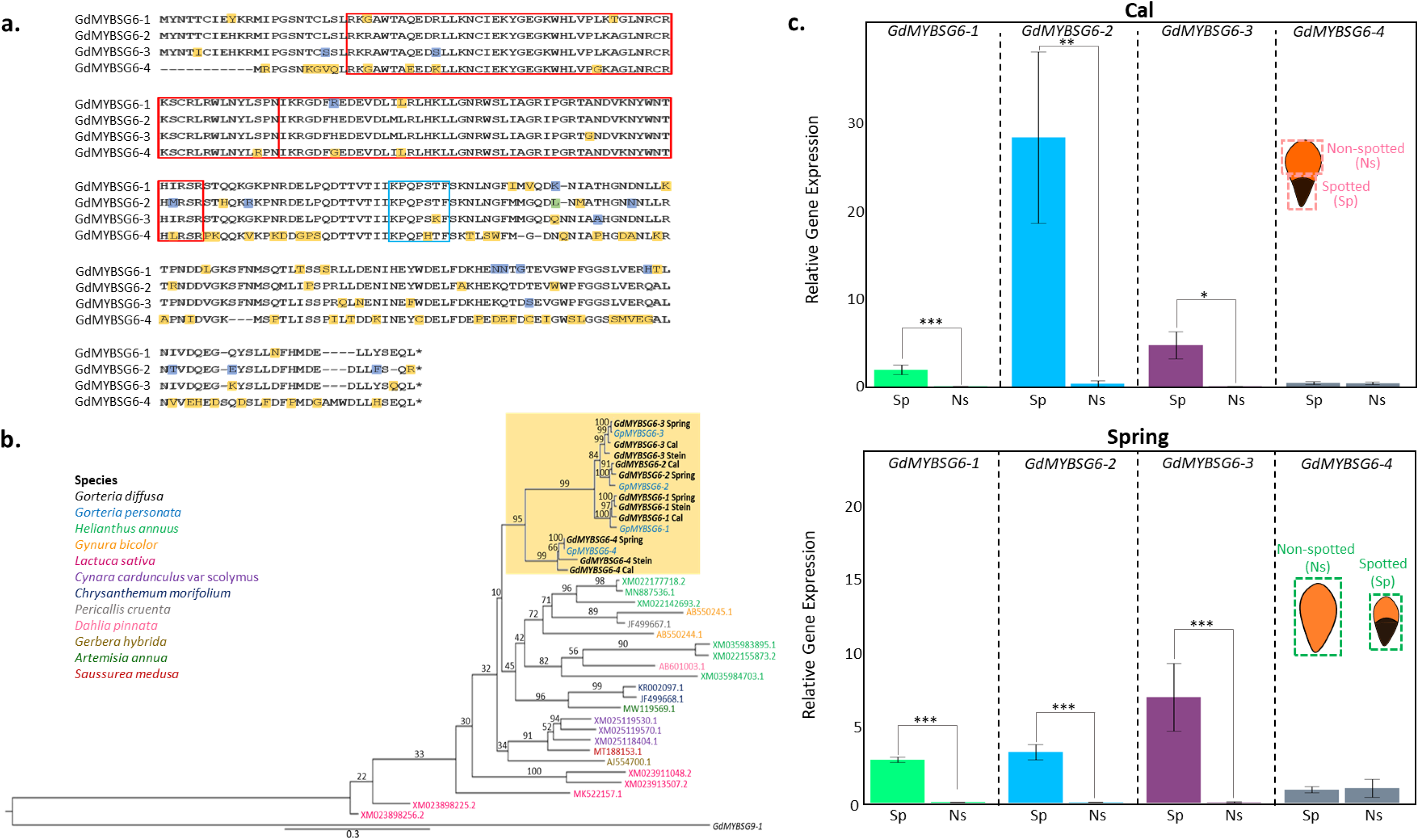
a) Protein alignment of GdMYBSG6 paralogues from the *G. diffusa* Spring morphotype. The red boxes indicate the R2 and R3 domains, and the blue box marks the subgroup 6 motif. b) Maximum likelihood phylogeny of Asteraceae subgroup 6 R2R3 MYB cDNA sequences. The tree was rooted with *GdMYBSG9-1*, a gene encoding a *G. diffusa* subgroup 9 R2R3 MYB used as the outgroup (Thomas 2009). Bootstrap values are given above each branch. *Gorteria* MYBSG6 genes sequenced during this project are indicated by the yellow box. Accession numbers are given, and corresponding species are colour coded as indicated in the key. c) qRT-PCR results showing the relative expression levels of *GdMYBSG6-1/2/3/4* in *G. diffusa* morphotypes Cal (top) and Spring (bottom). Cal spotted (Sp) and non-spotted (Ns) ray floret petal tissue and Spring whole spotted ray floret petals (Sp) and whole non-spotted ray floret petals (Ns) were sampled when spot formation initiates, as a small patch of pigment becomes visible. Error bars represent the mean ± s. e. of three biological replicates. Gene expression is relative to the reference gene *G. diffusa Elongation Factor 2* (*GdEF-2*). *p < 0.002, **p < 0.001, ***p < 0.0001.

### Phylogenetic analysis confirms that *GdMYBSG6-1 – 4* encode subgroup 6 R2R3-MYBs

A maximum likelihood phylogenetic reconstruction of the coding sequences of Asteraceae subgroup 6 R2R3-MYB transcription factors was used to assess the placement of *GdMYBSG6-1* – *4* within an evolutionary context (Figure 3b). The analysis confirmed that *Gorteria MYBSG6* genes clustered within subgroup 6, forming a single clade with high bootstrap support (95%). *G. personata MYBSG6* homologue sequences clustered with the corresponding *GdMYBSG6* orthologues, indicating that paralogues were already present prior to the speciation event that led to *G. diffusa* and *G. personata* emergence. Within the *Gorteria* clade, *MYBSG6-4* diverged first (bootstrap value 99%). The remaining *MYBSG6* genes formed a clade with *MYBSG6-2* and *MYBSG6-3* sister to one another (bootstrap value 84%). The Stein *MYBSG6-2* gene was excluded from the phylogeny because only a small section of the gene was successfully isolated.

### Three *GdMYBSG6* paralogues are upregulated in developing petal spots

To identify which, if any, of the *GdMYBSG6* genes were most likely to regulate pigment production during spot formation, the expression levels of each *GdMYBSG6* gene in spotted and non-spotted ray floret petal tissue were quantified across the three morphotypes at two landmarks of spot development (Figure S5): during spot initiation (Figure 3c), when a small patch of dark pigment is just visible, and later (Figure S6, Figure S7) as spot anthocyanin accumulates and specialised cell types emerge.

During spot initiation, *GdMYBSG6-4* had low expression in all morphotypes and similar expression levels in spotted and non-spotted petal tissue (Figure 3c). At the second developmental stage the expression of *GdMYBSG6-4* significantly increased in both spotted (t = 10.92 p < 0.001) and non-spotted (t = 6.84, p < 0.001) petal tissues of Cal (Figure S7), and in Stein (t = 2.94, p = 0.042) (Figure S6). Higher levels of Spring *GdMYBSG6-4* transcripts were also detected at this stage, but this upregulation was not statistically significant (Figure S7). Taken together these results indicate that *GdMYBSG6-4* expression patterns do not correlate with pigment accumulation in all three morphotypes, suggesting that this paralogue is unlikely to regulate anthocyanin production during spot formation.

In contrast, high expression levels of *GdMYBSG6-1*, *GdMYBSG6-2*, and *GdMYBSG6-3* were recorded in spotted tissue compared to non-spotted counterparts in both Cal (*GdMYBSG6-1*: t = 7.54, p < 0.001, 137-fold difference; *GdMYBSG6-2*: t = 6.35, p = 0.001, 84-fold difference; *GdMYBSG6-3*: t = 5.56, p = 0.002, 292-fold difference) and Spring (*GdMYBSG6-1*: t = 19.29, p < 0.001, 92-fold difference; *GdMYBSG6-2*: t = 13.91, p < 0.001, 154-fold difference; *GdMYBSG6-3*: t = 8.06, p < 0.001, 268-fold difference). Expression levels of *GdMYBSG6-2* in Cal spotted tissue were significantly higher than those of *GdMYBSG6-1* (t = 5.66, p < 0.001) and *GdMYBSG6-3* (t = 4.26, p < 0.001), while *GdMYBSG6-3* had the greatest expression levels in the Spring morphotype (*GdMYBSG6-1*: t = 2.97, p = 0.009; *GdMYBSG6- 2*: t = 2.49, p = 0.027) (Figure 3c). The expression of *GdMYBSG6-1*, *GdMYBSG6-2*, and *GdMYBSG6-3* in Spring and Cal morphotypes was extremely low or non-detectable in non-spotted petal tissue. Consistent with this, *GdMYBSG6-1* and *GdMYBSG6-3* expression levels were also very low in non-spotted ray florets from the Stein morphotype (Figure S6). At the later stage, *GdMYBSG6-1* and *GdMYBSG6-3* remained preferentially expressed in spotted tissues of Cal and Spring, and in Spring their expression levels decreased. The expression level of *GdMYBSG6-2* in spotted tissues, however, increased significantly at the second developmental stage in both Spring and Cal (Figure S7). These results indicate that the three paralogues could act redundantly to promote anthocyanin production during spot formation, with GdMYBSG6-2 likely to be the main driver.

### GdMYBSG6-1, 2 and 3 can induce anthocyanin production in *Nicotiana tabacum*

Stable transformations of tobacco plants (*N. tabacum*) were used to compare the biochemical properties of GdMYBSG6 proteins and assess their ability to induce anthocyanin synthesis in a heterologous context. The *GdMYBSG6-1*, *GdMYBSG6-2*, and *GdMYBSG6-3* genes that were upregulated in the spotted regions of *G. diffusa* petal tissue were placed under the control of strong constitutive 35S promoters and were each introduced into *N. tabacum* to generate stably transformed lines. Genotypic and phenotypic data were collected from the T_1_ generation corresponding to several (≥4) independent primary transformants for each gene.

Constitutive expression of *GdMYBSG6-1*, *GdMYBSG6-2*, or *GdMYBSG6-3* was sufficient to induce anthocyanin synthesis within all lines transformed with each construct. Petal tissue of all transformants had stronger anthocyanin phenotypes than wild type plants (Figure 4a). There was variation in the strength of the anthocyanin phenotype between lines within each set of *GdMYBSG6* transformants, but anthocyanin was ectopically produced within sepals, anthers, and leaves in at least a single line from each of *GdMYBSG6-1*, *GdMYBSG6-2*, and *GdMYBSG6-3* transformants (Figure 4a, Figure S10). Anthocyanin was extracted from anthers, sepals, and petals of transformants and wild type plants (Figure 4b, Figure S11). Multiple lines from each transformant and a minimum of two flowers from each plant, representing one line, were used in the analysis. Plants overexpressing any of these three *GdMYBSG6* had significantly higher anthocyanin concentrations in petals (*35S::GdMYBSG6-1*: t = 2.82, p = 0.03; *35S::GdMYBSG6-2*: t = 4.58, p = 0.002; *35S::GdMYBSG6-3*: t = 3.74, p = 0.006) and anthers (*35S::GdMYBSG6-1*: t = 2.40, p = 0.048; *35S::GdMYBSG6-2*: t = 5.83, p < 0.001; *35S::GdMYBSG6-3*: t = 3.58, p = 0.010) than wild type plants. Sepal anthocyanin content was also greater in *35S::GdMYBSG6-2* (t = 6.91, p < 0.001) and *35S::GdMYBSG6-3* (t = 2.73, p = 0.023) individuals compared to wild type (Figure S11). A single petal sample from each transformant was sent for UHPLC-MS/MS along with a wild type sample, no differences in anthocyanin composition were found (Figure S12, Figure S13).

**Figure 4.**
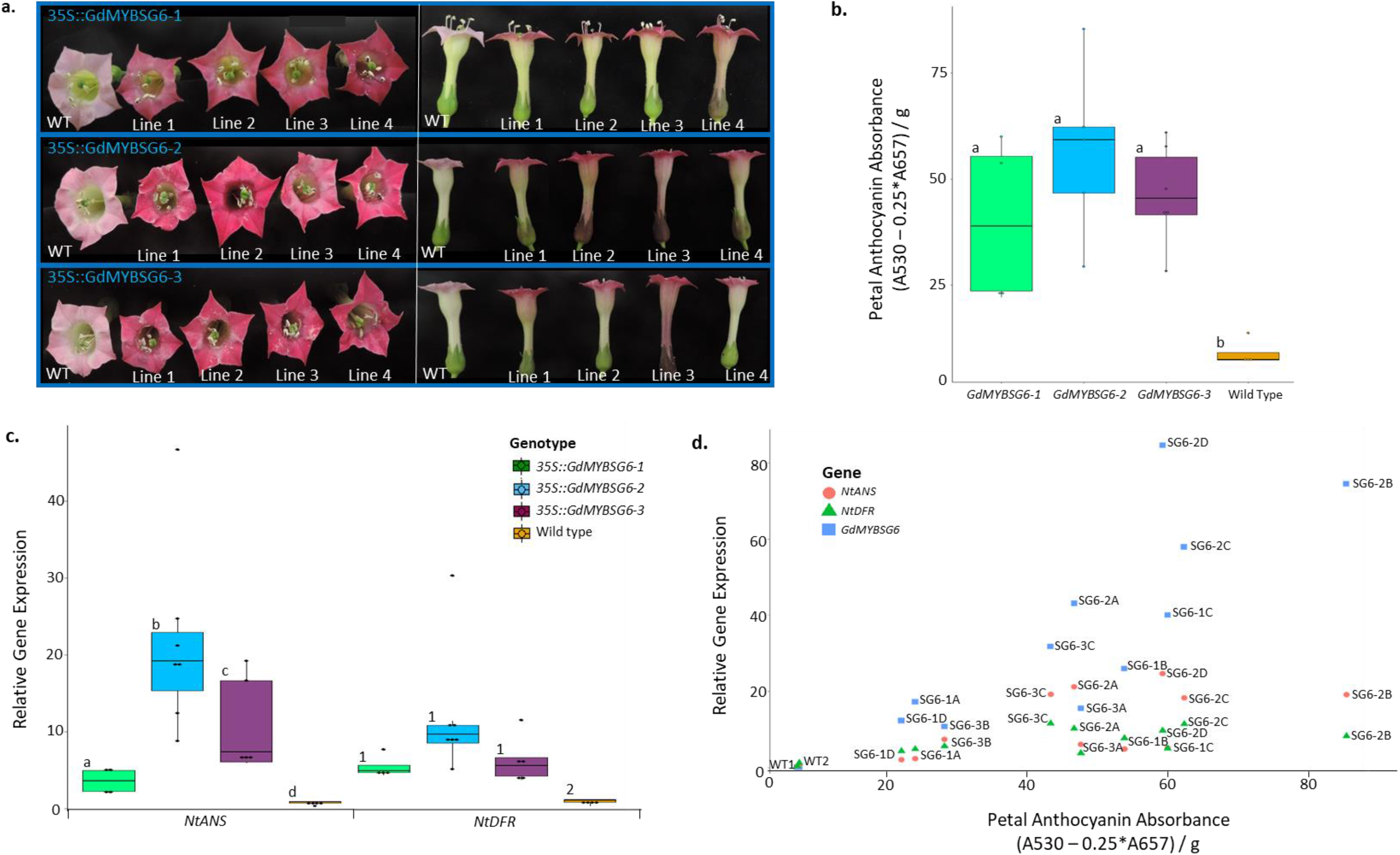
Anthocyanin phenotypes and gene expression levels in T1 *Nicotiana tabacum* plants stably transformed with either *GdMYBSG6-1, GdMYBSG6-2*, or *GdMYBSG6-3* under control of a strong constitutive promoter (35S). a) Flowers from several independent lines carrying each construct. Wild type flowers are labelled WT. b) Anthocyanin absorbance within petals. Each data point (black circle) is from a different independent line and represents the mean anthocyanin content of 4 – 6 samples. c) Relative expression levels of the genes encoding the anthocyanin synthesis enzymes anthocyanidin synthase (ANS) and dihydroflavonol 4-reductase (DFR) in the petals of *N. tabacum* transformants and wild type plants. Each data point (black circle) is from a different independent line and sample size is n = 4 – 7. d) Correlation between petal anthocyanin content and expression levels of genes of interest (*NtANS*, *NtDFR*, *GdMYBSG6- 1/2/3*) in individual plants from a subset of *N. tabacum* lines carrying each *35S::GdMYBSG6* construct. The line and construct are listed next to each data point (e.g. *35S::GdMYBSG6-1* line A = ‘SG6-1A’). qRT-PCR expression levels are colour coded according to which gene they correspond to (see key). Within a graph, tissues with statistically significant differences (p ≤ 0.05) in expression levels do not share a letter/number.

Late Biosynthesis Genes (LBGs) of the anthocyanin pathway represent potential downstream targets of GdMYBSG6 proteins. Therefore, the expression levels of two LBGs, *NtANS* and *NtDFR*, were investigated through qRT-PCR on multiple transgenic lines to determine whether expression patterns were consistent with transcriptional activation by GdMYBSG6 proteins (*35S::GdMYBSG6-1* n = 4, *35S::GdMYBSG6-2* n = 7, *35S::GdMYBSG6-3* n = 5). Transgene expression was generally higher across *35S::GdMYBSG6-2* lines compared to *35S::GdMYBSG6-1* and *35S::GdMYBSG6-3* lines (Figure S14). There was significant upregulation of *NtANS* (*35S::GdMYBSG6-1*: t = 2.37, p = 0.03; *35S::GdMYBSG6- 2*: t = 7.77, p < 0.001; *35S::GdMYBSG6-3*: t = 5.13, p < 0.001) and *NtDFR* (*35S::GdMYBSG6-1*: t = 5.39, p < 0.001; *35S::GdMYBSG6-2*: t = 8.25, p < 0.001; *35S::GdMYBSG6-3*: t = 5.96, p < 0.001) compared to wild type *N. tabacum*. *GdMYBSG6* expression and the expression of *NtANS* (t = 7.55, p = 0.001, R^2^ = 0.77) and *NtDFR* (t = 4.28, p < 0.001, R^2^ = 0.50) were positively correlated across transgenic lines.

The upregulation of *NtANS* in *35S::GdMYBSG6-1* transgenic lines was significantly lower than in *35S::GdMYBSG6-2* (t = 4.76, p = 0.001) and *35S::GdMYBSG6-3* (t = 2.50, p = 0.040) transformants, even when variability resulting from transgene expression levels was accounted for in the statistical model. In addition, *NtANS* expression was lower in *35S::GdMYBSG6-3* lines compared to *35S::GdMYBSG6-2* lines (t = 2.23, p = 0.044). A subset of *N. tabacum* transformant lines are used in Figure 4d to demonstrate that, despite similarities between certain lines regarding transgene expression levels and anthocyanin content, transgenic lines overexpressing *GdMYBSG6-1* had lower *NtANS* expression levels than *35S::GdMYBSG6-2* and *35S::GdMYBSG6-3* lines. These observations indicate that the three *GdMYBSG6* paralogues encode proteins with similar but not identical biochemical properties, and further comparisons between proteins would be required to address this comprehensively.

### *G. diffusa* genes encoding anthocyanin synthesis enzymes are upregulated in developing petal spots

To test whether genes encoding anthocyanin synthesis enzymes were also regulatory targets of GdMYBSG6 proteins in *Gorteria*, homologues of *ANTHOCYANIDIN SYNTHASE* (*GdANS*) and *DIHYDROFLAVONOL 4-REDUCTASE* (*GdDFR*) were isolated in *G. diffusa*. A single copy of *GdANS* was found and multiple *GdDFR* genes were identified with high sequence conservation. It remains unclear how many *GdDFR* genes are present, but our data suggest that at least two homologues exist. In addition, a malonyl transferase gene (*GdMAT1*) was investigated as another potential downstream target. Malonyl transferase (MAT) is an enzyme that catalyses the addition of malonyl groups to anthocyanins (Nakayama et al. 2003). As a high proportion of *G. diffusa* petal spot anthocyanins are malonylated, while other petal anthocyanins generally do not contain malonyl residues (Figure 2), a spot specific *GdMAT* could participate in spot elaboration. Four *G. diffusa* malonyl transferase variants were identified in a recent RNA-seq analysis (Kellenberger unpublished). All contained the motif associated with the subfamily for anthocyanin malonyl transferases (Unno et al. 2007). Expression levels were much higher within spotted petal tissue than non-spotted tissue for one of these variants, *GdMAT1* (Kellenberger unpublished).

If *GdANS*, *GdDFR*, and *GdMAT1* are regulatory targets of GdMYBSG6, their expression profiles are likely to follow the expression pattern of GdMYBSG6s. To test this hypothesis, we characterised their expression profiles in the same tissue samples as those used to establish the transcriptional dynamics of *GdMYBSG6* genes. Expression analyses were designed to detect all *GdDFR* transcripts that contained the catalytic triad of amino acids (Dodson and Wlodawer 1998; Petit et al. 2007). As spot formation initiates, *GdANS*, *GdDFR*, and *GdMAT1* were found to be significantly upregulated in spotted compared to non-spotted petal tissue in Cal (*GdANS*: t = 4.62, p = 0.002; *GdDFR*: t = 7.64, p < 0.001, *GdMAT1*: t = 4.94, p = 0.003) and Spring (*GdANS*: t = 7.07, p < 0.001; *GdDFR*: t = 13.25, p < 0.001, *GdMAT1*: t = 5.24, p = 0.002) (Figure 5a). At the later stage, the expression levels of both *GdANS* and *GdDFR* significantly increased in non-spotted petal tissue of Cal (*GdANS*: t = 10.67, p < 0.001; *GdDFR* t = 23.6, p < 0.001) and Spring (*GdANS*: t = 10.86, p < 0.0001; *GdDFR*: t = 17.15, p < 0.0001) morphotypes, coinciding with the production of anthocyanin pigmentation on the abaxial side of the ray floret petals at this stage (Figure S5, Figure S15). At this later developmental stage there was no difference in expression levels of *GdANS* between the spotted and non-spotted tissues. However, *GdDFR* remained upregulated in the spotted region of both Cal (t = 5.56, p = 0.002) and Spring (t = 2.79, p = 0.024). *GdMAT1* was also upregulated at the later developmental stage in both Cal (t = 3.80, p = 0.01) and Spring (t = 4.31, p = 0.005), with expression in non-spotted tissue remaining very low (Figure S15). In the non-spotted Stein petals *GdMAT1* expression was also very low and non-detectable for some biological replicates (mean ± s.e: dev 1 0.01 ± 0.004, dev 2 0.012 ± 0.01).

**Figure 5.**
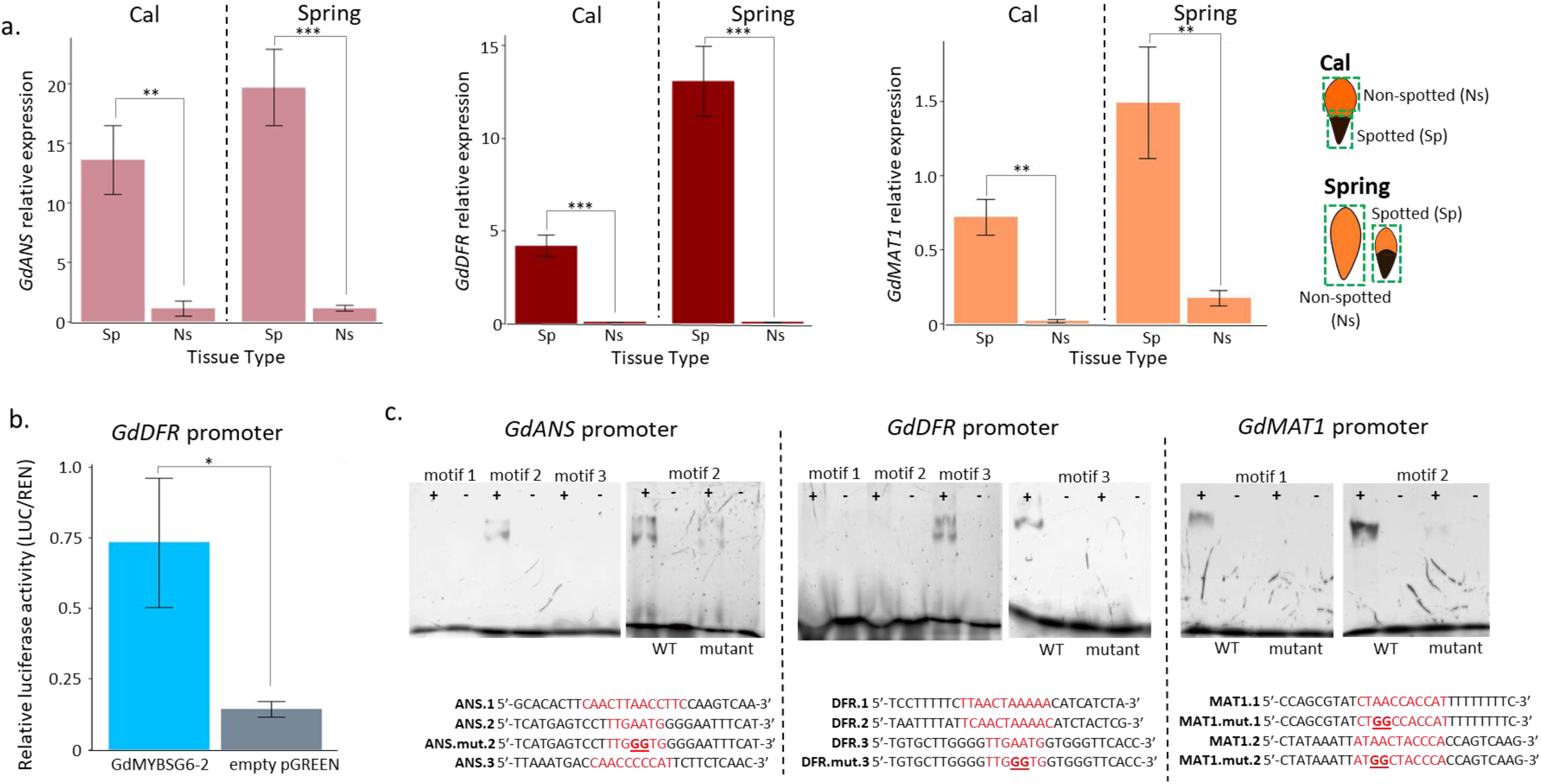
GdMYBSG6 proteins are likely transcriptional regulators of genes encoding late anthocyanin synthesis enzymes in *G. diffusa*. a) Relative expression of *G. diffusa ANTHOCYANIN SYNTHASE* (*GdANS*), *DIHYDROFLAVONOL 4-REDUCTASE* (*GdDFR*) and *MALONYL TRANSFERASE 1* (*GdMAT1*) in spotted (Sp) and non-spotted (Ns) ray floret petal tissue sampled from Cal and Spring morphotypes as the spot initiates and just becomes visible as a small patch of pigment. b) Relative luciferase activity indicating whether GdMYBSG6-2 can activate luciferase expression driven by the *GdDFR* promoter region (*GdDFRp*), with the empty effector plasmid (pGreen) used as a negative control. c) EMSAs testing whether GdMYBSG6-2 can bind to wild-type and mutated versions of selected DNA motifs from *GdANSp*, *GdDFRp*, and *GdMAT1p*. Negative controls (-) consist of binding reactions with no protein added. DNA motif sequences are given under each gel, with predicted binding sites highlighted in red and changes made in mutated motifs indicated in underlined bold. *p < 0.05 ** p < 0.002 ***p < 0.0001.

### GdMYBSG6-1 - 3 can bind to promoter regions of *GdDFR*, *GdANS*, and *GdMAT1*

To test whether GdMYBSG6-1 – 3 could act as direct activators of the selected LBGs, we tested the ability of a recombinant version of GdMYBSG6-2 to bind *in vitro* to the potential promoter regions of *GdDFR*, *GdANS* and *GdMAT1*. First, regions immediately upstream of the start codon of these anthocyanin biosynthesis enzymes were isolated (*GdMAT1p:* 1016 bp upstream of *GdMAT1* coding sequence, *GdANSp:* 334 bp upstream of *GdANS* coding sequence). Multiple promoter sequences were found for *GdDFR*, likely because multiple homologues are present within the *G. diffusa* genome. All of these *GdDFR* upstream variants showed high sequence conservation from approximately 380 bp upstream of the start codon. As such, this 380 bp common *GdDFRp* region was used for subsequent experiments. We identified potential MYB binding sites within these upstream regions using consensus sequences reported in previous studies (see Methods) and then used EMSAs to determine whether GdMYBSG6-2 could bind to these motifs. Heterologous expression in tobacco suggests that GdMYSG6-1, 2 and 3 have similar properties and GdMYBSG6-2 was chosen for this binding assay as our expression data singled it out as the top candidate to regulate anthocyanin synthesis during spot development. EMSA results demonstrated that GdMYBSG6-2 was able to bind specific DNA motifs within the upstream regions of *GdANS*, *GdDFR*, and *GdMAT1* (Figure 5c). Substituting ‘AA’ with ‘GG’ in the centre of those motifs was sufficient to reduce or abolish the binding capacity of GdMYBSG6-2 in all cases. Both motifs tested from *GdMAT1p* were bound by GdMYBSG6-2. Out of three motifs tested in each of *GdANSp* and *GdDFRp*, GdMYBSG6-2 consistently bound to only one motif in each upstream region. These bound motifs contained the sequence TTGAATG – previously identified by Wang et al. (2013) in the *DFR2* promoter of *Gerbera*. Next, we tested whether GdMYBSG6-2 was able to activate the transcription of *GdDFR* by dual-luciferase assay in *N. benthamiana*. We found that the GdMYBSG6-2 protein induced the expression of luciferase driven by the *GdDFR* promoter relative to the control (t = 3.30, p = 0.028) (Figure 5b). Taken together, these results indicate that GdMYBSG6-2 can bind to promoter regions of *GdANS*, *GdMAT1*, and *GdDFR* and activate gene expression driven by the *GdDFR* promoter in a heterologous system.

## Discussion

*Gorteria diffusa* is a daisy species inhabiting a narrow endemic range within the Succulent Karoo biodiversity hotspot of South Africa. High levels of geographically defined floral variation are present within *G. diffusa*, making this non-model species an excellent system to explore the evolution of natural diversity. Key components of this intraspecific diversity are unusually elaborate petal spots, several of which can mediate sexual deception of the bee-fly that pollinates *G. diffusa*. Richly coloured and deeply textured, these spots are composed of several specialised cell types that create tridimensional elaborations and varying colours. Here we investigated one aspect, pigment production, of the processes that underpin complex spot formation. We found that malonylated cyanidin 3-glucoside almost exclusively accumulates in *G. diffusa* petal spots. Our study further identifies three paralogous *GdMYBSG6* genes likely to activate spot specific anthocyanin production through transcriptional control of *GdDFR*, *GdANS,* and *GdMAT1* - three genes encoding late anthocyanin biosynthetic enzymes in developing petal spots.

### Petal anthocyanin composition is similar between morphotypes, but spot anthocyanin concentrations differ

Differences in anthocyanin composition between cultivars have been reported in Asteraceae species including *Dahlia variabilis* and *Gerbera jamesonii* (Takeda et al. 1986) but this was not the case between the three *G. diffusa* morphotypes we examined. Anthocyanin composition was largely consistent across the morphotypes; cyanidin 3-glucoside was the predominant anthocyanin and all anthocyanins identified were derived from cyanidin (with a single unconfirmed exception). Cyanidin derivatives also contribute toward pink to red floral colouration in *Chrysanthemum morifolium* and *Lilium* spp. and blue colouration in *Meconopsis grandis* and *Centaurea cyanus* (Hong et al. 2015; Suzuki et al. 2016; Yoshida et al. 2006; Yoshida and Negishi 2013). One exception to this consistency between *G. diffusa* morphotypes, was that all petal tissues from Cal lacked anthocyanins containing caffeate, which were present in small quantities in Stein non-spotted ray florets, Spring petal spots, and in the region with ‘marks’ on Spring non-spotted ray florets.

Spring petal spots contained significantly more anthocyanin than the petal spots of Cal. This may be partly due to differences in spot micromorphology, in particular papillae: the groups of swollen epidermal cells. Papillae are black and swollen in the Spring morphotype; whereas, in Cal, papillae are often dark green in colour, smaller, and are more sparsely distributed. Cal spots have a raised appearance largely due to curvature of the ray floret petals as opposed to enlarged pigment epidermal cells, while Spring spotted ray florets exhibit less curvature. Anthocyanins may be less dominant in Cal petal spot pigmentation, with other pigments potentially contributing toward the dark green colouration. Preliminary investigations suggest that chlorophyll concentrations could be significantly higher in Cal compared to Spring petal spots (Walker, 2012).

### Anthocyanin malonylation is a characteristic of petal spot pigments

We found that most anthocyanins within *G. diffusa* petal spots (approximately 60%) are acylated by malonate, while only small quantities of malonylated anthocyanins are present in non-spotted petal regions. This difference is highly consistent across the Cal, Stein, and Spring morphotypes. Malonic acid is the most frequent aliphatic acyl group in acylated anthocyanins and is found throughout the Asteraceae, including in *Senecio cruentus* and *Gerbera* (Harborne 1963; Takeda et al. 1986). The specific role of malonylated anthocyanins in pigmenting *G. diffusa* petal spots is unknown, but studies in *Dahlia* and *Arabidopsis* suggest that malonic acid acylation could increase the stability of anthocyanins (e.g. Luo et al. 2007, Suzuki et al. 2002). One way this is achieved is through the formation of anthocyanin zwitterions; protons disassociate, and this can decrease the pH of vacuolar sap protecting the anthocyanin from degradation induced by any pH increases (Takeda et al. 1986, Figueiredo et al. 1999). Anthocyanins which are acylated have also been reported to be more stable against heat and light stress (Inami et al. 1996; Sadilova et al. 2006; Xu et al. 2017; Zhao et al. 2017).

*G. diffusa* grow within a desert and flowers open from mid-morning to mid-afternoon, so the adaxial petal surface is exposed to high light intensities and may experience increased temperatures for a prolonged period. As such, having malonylated anthocyanins within the spot could protect against anthocyanin degradation resulting from light exposure. This may be particularly important for petal spots because they have a key role in attracting pollinators (Ellis and Johnson 2010; Johnson and Midgley 1997). Floral temperatures can also play a role in attracting pollinators, with flowers providing a heat reward in some species (Creux et al. 2021; Harrap et al. 2017; Van Der Kooi et al. 2019; Sapir et al. 2006). If this occurs within *G. diffusa* floral heating may be advantageous, with high anthocyanin stability in petal spots mitigating against the associated detrimental consequences of anthocyanin degradation. In addition, heat and light stress may be less severe for abaxial pigmentation as abaxial petal surfaces are only exposed to direct sunlight when the flowers are closed in the early morning and late afternoon.

Anthocyanin composition is strongly shifted in favour of anthocyanin malonylation in both Cal and Spring petal spots. However, in Spring, qualitative modification of pigment identity is also accompanied by an increase in overall pigment production in the spot. This implies that spot development in the Spring morphotype relies on two key events: a change in anthocyanin modification profile (as observed in Cal) and an overall increase in pigment production compared to non-spotted ray florets. Interestingly, the shift to higher proportions of malonylated anthocyanins also occurred within the ‘marks’ of Spring. These patches of black pigment are not accompanied by any other form of cellular elaborations (Figure S1) and so this distinct spot anthocyanin composition is not unique to complex spot phenotypes but occurs in all anthocyanin patterning on the adaxial petal surface. Taken together these results suggest that anthocyanin malonylation might be one of the earliest events in spot specification.

### *GdMYBSG6-1* – *3* encode transcriptional activators of petal spot anthocyanin production

We isolated four genes encoding subgroup 6 R2R3-MYB transcription factors that are expressed within *G. diffusa* ray floret petals. *GdMYBSG6-1, 2* and *3* were shown to be likely regulators of spot anthocyanin production. These paralogues are strongly upregulated within spotted petal tissue during spot development and can trigger ectopic anthocyanin production when constitutively overexpressed in a tobacco heterologous context. In addition, gel-shift assays demonstrated that the GdMYBSG6-2 protein, chosen as our representative paralogue of GdMYBSG6-1, 2 and 3 behaviour, could bind to motifs within promoter regions of *GdANS*, *GdDFR*, and *GdMAT1*. All of these genes are potential downstream targets of GdMYBSG6s and display expression patterns consistent with regulation by *GdMYBSG6-1 – 3*. Evidently, *G. diffusa* anthocyanin petal patterning results from the spatially restricted transcription of MYB gene activators. This is consistent with findings in other systems including *Lilium* (Yamagishi et al. 2014), *Mimulus* (Ding et al. 2020; Yuan et al. 2014), and *Clarkia* (Martins et al. 2017). This transcriptional regulation is typically mediated by an MBW complex which contains a basic helix-loop-helix (bHLH), a WD-repeat (WDR), and a MYB protein; the latter is thought to be the main driver of DNA binding specificity, directly impacting the developmental role of the complex (Ramsay and Glover 2005). *GdMYBSG6-4* had equivalent expression levels in spotted and non-spotted petal regions and its upregulation coincided with the production of abaxial petal pigmentation in non-spotted petal tissue. Further characterisation is required to establish *GdMYBSG6- 4* function and *in situ* hybridisation could be informative to establish whether *GdMYBSG6-4* transcripts are localised to the abaxial petal surface.

### Differences in *Gorteria MYBSG6* expression patterns mirror evolutionary divergence

Phylogenetic analysis confirmed that the four paralogous *Gorteria MYBSG6* genes form a clade within subgroup 6 of the R2R3-MYB family. *G. diffusa* and *G. personata* representatives cluster according to each paralogue indicating that the gene duplication that led to the formation of this clade predates the speciation event. The first gene to diverge within the phylogeny is *GdMYBSG6-4* which lacks spot-specific expression. The remaining *GdMYBSG6-1* – *3* genes, that are all highly upregulated in spotted petal tissue during spot development, form a clade. This implies that the common ancestor of these three genes likely had a spot-specific expression pattern. Signatures of positive selection in recently duplicated R2R3-MYBs have been reported in multiple MYB gene lineages (Jia et al. 2003). Functional divergence through sub- or neofunctionalization (e.g. Haberer et al. 2004) are two of several possible evolutionary outcomes following duplication. In monkeyflowers, new floral pigmentation patterns occurred following duplications of MYB anthocyanin regulators: *M. cupreus* and *M. luteus*. var *variegatus* evolved petal lobe anthocyanin in parallel following a duplication event within a different MYB locus in each species (Cooley et al. 2011). Whether spot-expression was gained along the lineage leading to *GdMYBSG6-1 – 3* or lost along the *GdMYBSG6-4* branch is not clear, but it is possible that gene duplications resulting in the expression of multiple *GdMYBSG6* genes within the *G. diffusa* petal spot contributed toward the evolution of such elaborate spot phenotypes. Perhaps, having three spot anthocyanin regulators is advantageous as it increases gene dosage leading to higher concentrations of GdMYBSG6 proteins to enhance anthocyanin production (Cheng et al. 2018, Kondrashov and Kondrashov 2006). *GdMYBSG6-2* and *GdMYBSG6-3* likely encode the dominant transcriptional activators of anthocyanin production in *G. diffusa* petal spots because they have the highest expression levels during spot initiation in both Cal and Spring petal spots. *GdMYBSG6-2* also had significantly elevated expression in both morphotypes relative to the other *GdMYBSG6* genes during the later developmental stage, when specialised cell types are differentiating. GdMYBSG6-1, 2 and 3 may have divergent roles in spot anthocyanin production if, for example, expression of one gene is localised to certain spot regions or is specific to a single specialised cell type. Future work could address this by investigating the expression of each paralogue at greater spatial resolution, comparing between different subregions of the developing spot. Comparisons between *N. tabacum* plants expressing each *GdMYBSG6* paralogue indicate potential differences in biochemical properties as the extent to which each protein upregulated *NtANS* overall and relative to *NtDFR* varied. While the divergence between *GdMYBSG6-4* and GdMYBSG6-1 *–* 3 involves a clear change in expression pattern, sub- or neofunctionalization between *GdMYBSG6-1*, *GdMYBSG6-2*, and *GdMYBSG6-3*, if present, could involve ongoing divergence in protein properties.

### GdMYBSG6-1, 2 and 3 likely regulate the expression of genes encoding the late anthocyanin synthesis enzymes *GdANS* and *GdDFR*

Individual subgroup 6 R2R3-MYB transcription factors can regulate multiple genes encoding anthocyanin synthesis enzymes (Petroni and Tonelli 2011). Here, we investigated the late biosynthesis genes, *GdANS* and *GdDFR*, as potential downstream targets of GdMYBSG6 proteins within the *G. diffusa* anthocyanin synthesis pathway. A single *GdANS* gene was identified and at least two *GdDFR* genes. The spatio-temporal expression of *GdANS* and combined *GdDFRs* largely correlated with the appearance of anthocyanin pigmentation within the ray florets. These genes were highly expressed in spotted petal tissue during spot initiation, with low expression levels in non-spotted petal tissue. During the later developmental stage both enzymes-encoding genes remained highly expressed in spotted tissue, allowing for continued anthocyanin production as specialised cell types emerged – including papillae which expand and fill with anthocyanin. At this later stage *GdANS* and *GdDFR* transcription also increased in non-spotted petal tissue as abaxial petal pigmentation appears. It is unclear whether the same *GdDFR* genes are upregulated across the entire ray floret petal or whether there is a spot specific *GdDFR* gene or allele. Multiple copies of floral anthocyanin synthesis enzymes have been reported in other systems including Asiatic hybrid lilies (*Lilium*) (Nakatsuka et al., 2003; Lai et al. 2012), and *Clarkia gracilis*. A *DFR* allele is expressed only in the spotted region of the *C. gracilis* petal and prior to other *DFRs* that synthesise background petal pigmentation (Martins et al. 2013). Further investigation is required to determine whether similar functional divergence exists between the *GdDFR* homologues potentially expressed within *G. diffusa* petals. However, all *GdDFR* promoter regions obtained to date have a highly conserved region containing the motif that GdMYBSG6-2 bound to. This suggests that all *GdDFR* homologues are regulated in the spot by GdMYBSG6, but other *cis*- elements may differ between homologues.

### GdMYBSG6-1, 2 and 3 likely regulate a malonyl transferase enzyme expressed within petal spots

As a high proportion of spot anthocyanins are malonylated we investigated whether a gene encoding a malonyl transferase was upregulated in petal spots. Malonyl transferases catalyse the addition of malonyl groups to anthocyanins. We found that *GdMAT1* was expressed at significantly higher levels in spotted compared to non-spotted petal tissue during spot initiation and elaboration in both Cal and Spring, while expression levels were very low in non-spotted Stein. These expression patterns mirror those of *GdMYBSG6-*1, 2 and 3, and gel shift assays confirmed that GdMYBSG6-2 can bind to the promoter region of *GdMAT1*. The regulation of malonyl transferases by MYB transcription factors has been investigated in few studies to date. Leaf transcriptomics followed by a gene regulatory network analysis predicted a likely interaction in *Brassica rapa* (Rameneni et al. 2020). Additionally, expression analyses conducted on berry tissues from vine plants (*Vitis vinifera*) overexpressing or silencing *VvMYBA* indicate that VvMYBA regulates the transcription of the acyltransferase *Vv3AT* (Rinaldo et al. 2015). Here, we identified malonylation as a key event in *G. diffusa* petal spot formation along with a potential transcriptional activator of this process. These components of the gene regulatory network controlling spot pigmentation may be under selection, contributing to the evolution of elaborated sexually deceptive spots in *Gorteria*.

## Conclusion

Through gene expression analyses and functional investigations, we have identified three recently duplicated subgroup 6 R2R3-MYB transcription factors (GdMYBSG6-1, 2 and 3) that could activate anthocyanin production during the development of elaborate petal spots in *G. diffusa*. We have demonstrated that the expression patterns of these paralogous genes are spatially restricted to the petal region in which spots emerge and we found genes encoding the anthocyanin synthesis enzymes GdDFR and GdANS are likely targets of these regulators. We discovered that anthocyanin malonylation is a characteristic of *G. diffusa* spot pigmentation and the GdMYBSG6 transcription factors we characterised also likely regulate the gene encoding the spot-specific malonyl transferase GdMAT1. Whether this is specific to the *Gorteria* genus or a general chemical trick plants use to produce salient motifs on their petals, and whether the addition of a malonyl group plays a role in the formation of a sexually-deceptive spot, represent exciting avenues for future investigations.

## Acknowledgements

We thank Matthew Dorling and Emma Jackson for excellent plant care; Boris Delahaie, Paula Rudall, Diarmuid Ó’Maoiléidigh, Julian Hibberd, and Emily Bailes for valuable discussions and feedback; Allan Ellis for supplying the seeds and sharing his expertise of the system; Amy Milburn, Daniel Canniffe, François Parcy, Michael Batie, and Sonia Roche for providing equipment and advice on completing EMSA experiments. We also thank Eugenio Butelli for sharing plasmids for the dual-luciferase assay, and Matthew Dorling and Boris Delahaie for their excellent photographs of Cal and Spring capitula, respectively. R.F. was supported by a University of Cambridge NERC Doctoral Training Programme grant (grant number NE/L002507/1) and G.M. was supported by a BBSRC-DTP doctoral fellowship. This study was also supported by a Swiss National Science Foundation (SNSF) Early Postdoc. Mobility grant to R.T.K. (fellowship P2ZHP3_178043), a fellowship from the Gatsby Charitable Foundation to E.M., an Isaac Newton Trust grant to R.T.K. and B.J.G., a Natural Environment Research Council (NERC) grant to B.J.G. (NE/P011764/1), a BBSRC grant to B.J.G. (BB/V000314/1) and a Cambridge Trusts Scholarship to F.K.

## Author Contribution

R.F., G.M., and B.J.G. designed the study. R.W. and R.K. provided transcriptome data; R.K. provided draft genome data; Q.W. performed the phylogenetic analysis; G.M. established a qRT-PCR reference gene; G.M., R.F. and F.K. gene hunted; G.M. and R.F. generated the plant expression vectors and transgenic tobacco; R.F. performed and analysed the qRT-PCR experiments; E.S., E.M. and R.F identified candidate binding motifs; R.F. and E.M. performed the EMSAs; R.F. genotyped, phenotyped, and imaged the tobacco transformants; F.K. performed the luciferase experiments with input from E.H.S.; L.H. performed the UHPLC-MS/MS analysis, identified the anthocyanins, and L.H. and R.F. analysed the data; R.F. prepped UHPLC-MS/MS samples and performed and analysed *G. diffusa* spectrophotometer anthocyanin quantifications; R.F. and F.K. took the images of dissected capitula; R.F. completed all statistical analyses; E.M., E.H.S. and B.J.G advised on experimental design; R.F. prepared the figures with input from E.M. and B.J.G; R.F. wrote the first draft of manuscript used by R.F., E.M. and B.J.G to prepare the subsequent versions. All other authors read and provided feedback on the final draft and approved the final version, with the exception of G.M. who is deceased but whose insight from project discussions was highly valued and actively taken into account upon creation of the manuscript.

