## Supplementary Information for "Complex petal spot formation in the Beetle Daisy (*Gorteria diffusa*) relies on spot-specific accumulation of malonylated anthocyanin regulated by paralogous GdMYBSG6 transcription factors"

The following Supporting Information is available for this article:

**Fig. S1** Anthocyanin pigmentation phenotypes in the Spring morphotype

**Fig. S2** Pigmentation on the abaxial sides of *G. diffusa* ray floret petals

**Fig. S3** Chromatograms from *G. diffusa* UHPLC-MS/MS

**Fig. S4** Mass spectra from *G. diffusa* UHPLC-MS/MS

**Fig. S5** *G. diffusa* ray floret developmental stages used in qRT-PCRs

**Fig. S6** Gene expression analysis in the non-spotted petals of the Stein morphotype

**Fig. S7** Gene expression analysis in the Cal and Spring morphotypes

**Fig. S8** Verification of transgene expression in transformed *N. tabacum* using RT-PCR

**Fig. S9** Developmental stages used for qRT-PCR and pigment extraction in *N. tabacum*

**Fig. S10** Leaves of *N. tabacum* transgenic lines constitutively overexpressing *GdMYBSG6* genes

**Fig. S11** Anthocyanin extractions from floral tissue of transgenic *N. tabacum* lines constitutively overexpressing *GdMYBSG6* genes

**Fig. S12** Chromatograms of anthocyanin petal extractions from transgenic *N. tabacum* constitutively overexpressing *GdMYBSG6* genes

**Fig. S13** Spectra of anthocyanin petal extractions from transgenic *N. tabacum* constitutively overexpressing *GdMYBSG6* genes

**Fig. S14** *GdMYBSG6* relative expression levels in floral tissue of transgenic *N. tabacum* carrying a *35S::GdMYBSG6* transgene

**Fig. S15** Expression analysis of *G. diffusa* genes coding for late anthocyanin synthesis enzymes

**Table S1** Primer sequences

**Table S2** Anthocyanin quantification and characterisation

**Table S3** Anthocyanins detected in the ray floret tissue of *G. diffusa*

**Methods S1** Anthocyanin quantification

**Methods S2** Map of plasmids used in *N. tabacum* transformation

**Methods S3** *N. tabacum* stable transformation procedure

**Methods S4** Map of plasmid used to produce the recombinant GdMYBSG6-2 protein used in EMSA

**Methods S5** Buffer solutions used for recombinant protein purification and EMSA

**Methods S6** Map of plasmid used in dual luciferase assays

**Fig. S1** Anthocyanin pigmentation phenotypes in the Spring morphotype

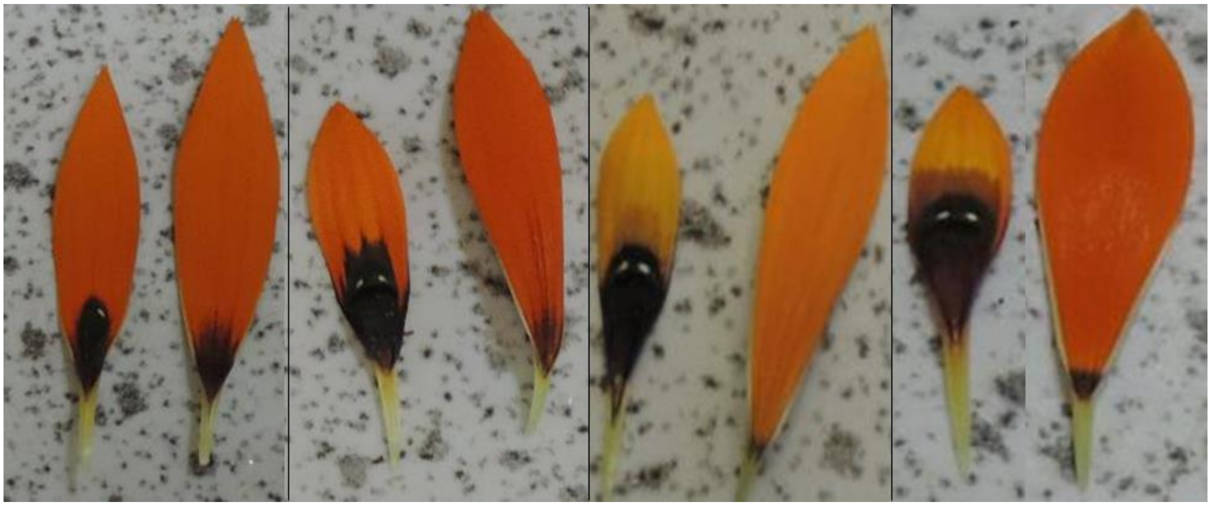

Spotted (left side of each panel) and non- spotted ray florets (right side of each panel) from different individuals encompassing the variation in Spring ray floret phenotypes. The spotted ray floret on the extreme left is only partially developed. The dark patch ‘mark’ at the base of the non-spotted ray floret varied in size between individuals and in some cases was completely absent.

**Fig. S2** Pigmentation on the abaxial sides of *G. diffusa* ray floret petals

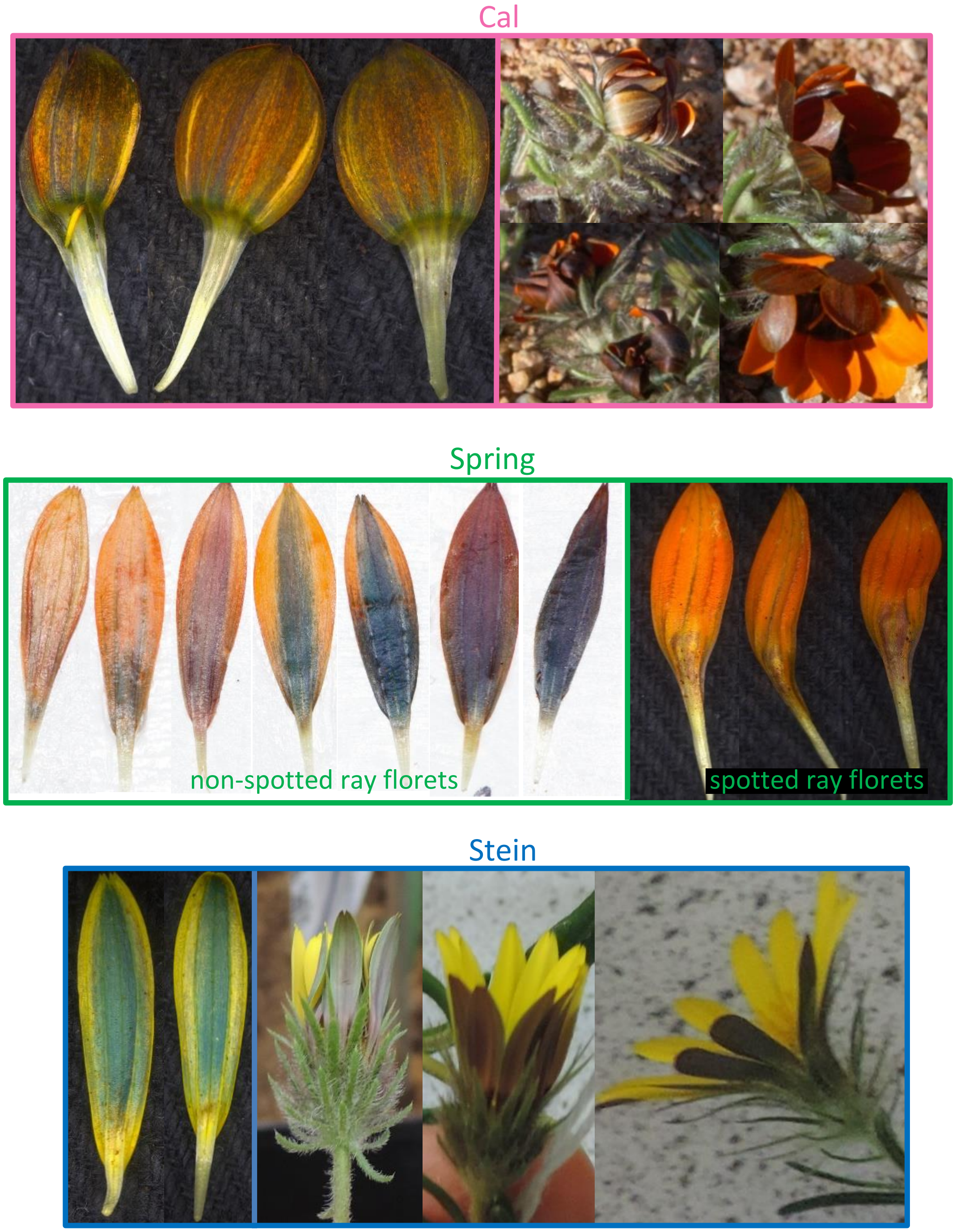


Photographs of the abaxial sides of *G. diffusa* ray florets in the morphotypes Cal, Spring, and Stein.

**Fig. S3** Chromatograms from *G. diffusa* UHPLC-MS/MS

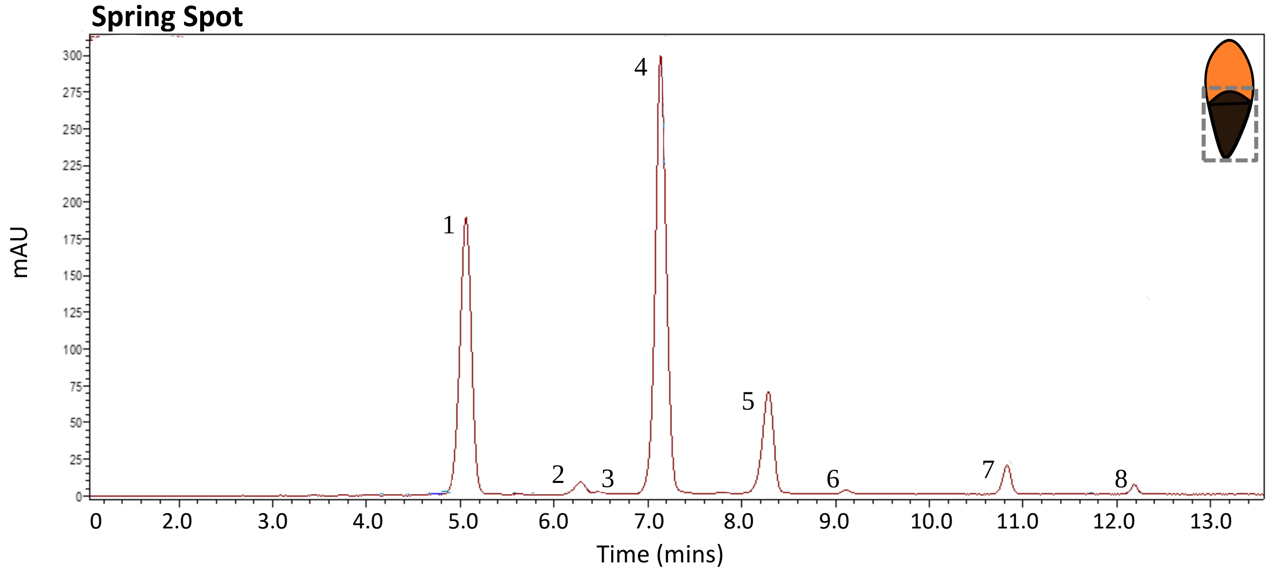


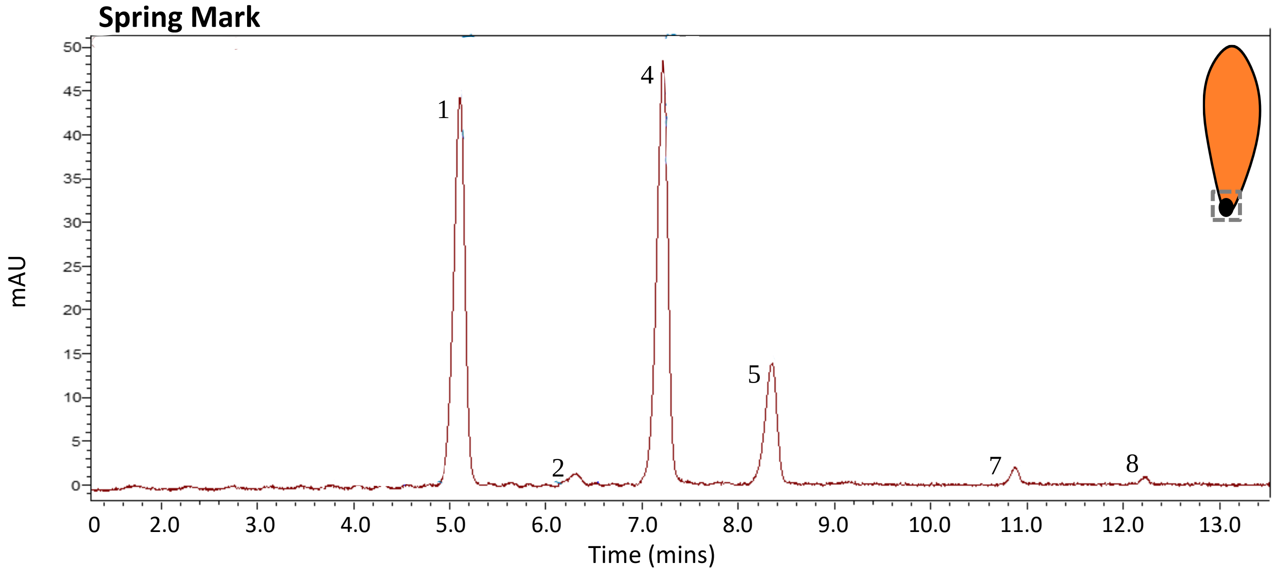


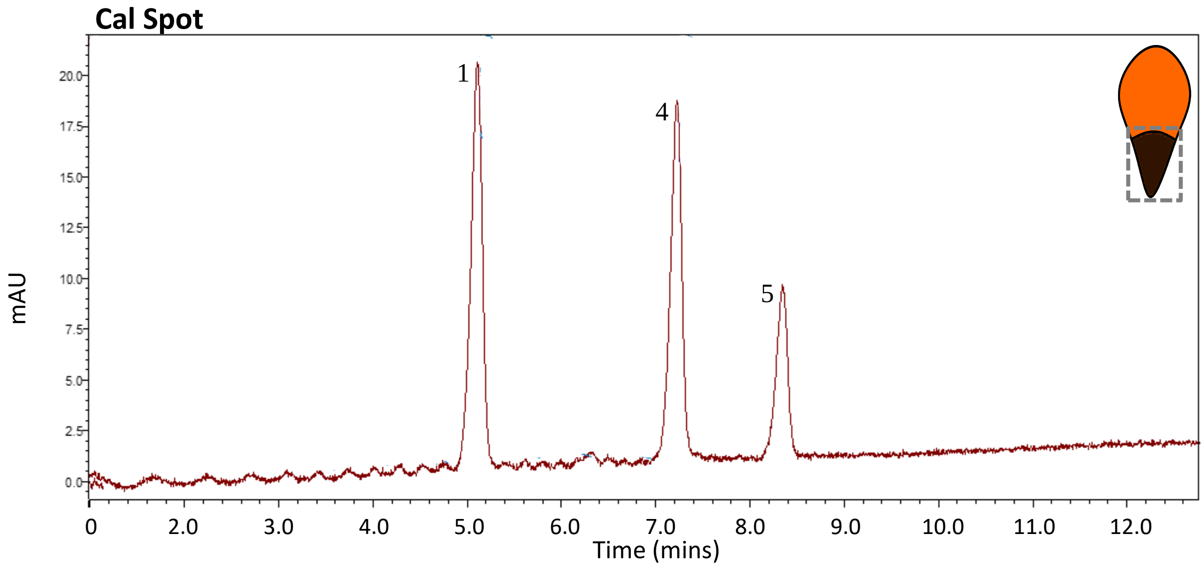

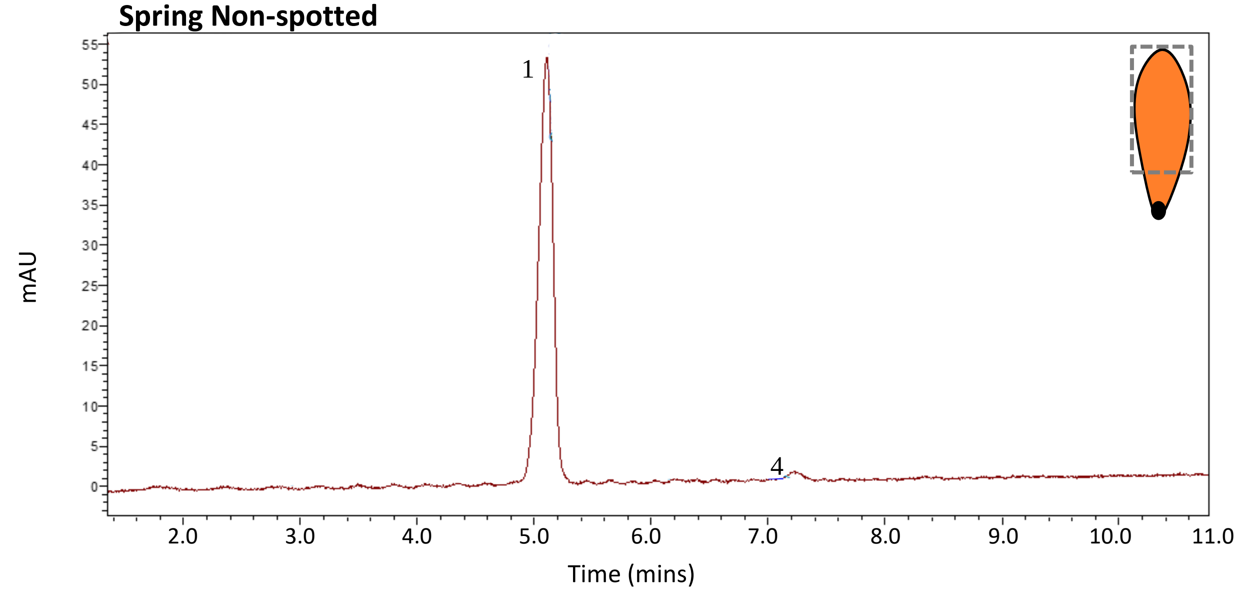
Spring Non-spotted
mAU
Time (mins)
1
4
0
5
10
15
20
25
30
35
40
45
50
55
2.0
3.0
4.0
5.0
6.0
7.0
8.0
9.0
10.0
11.0

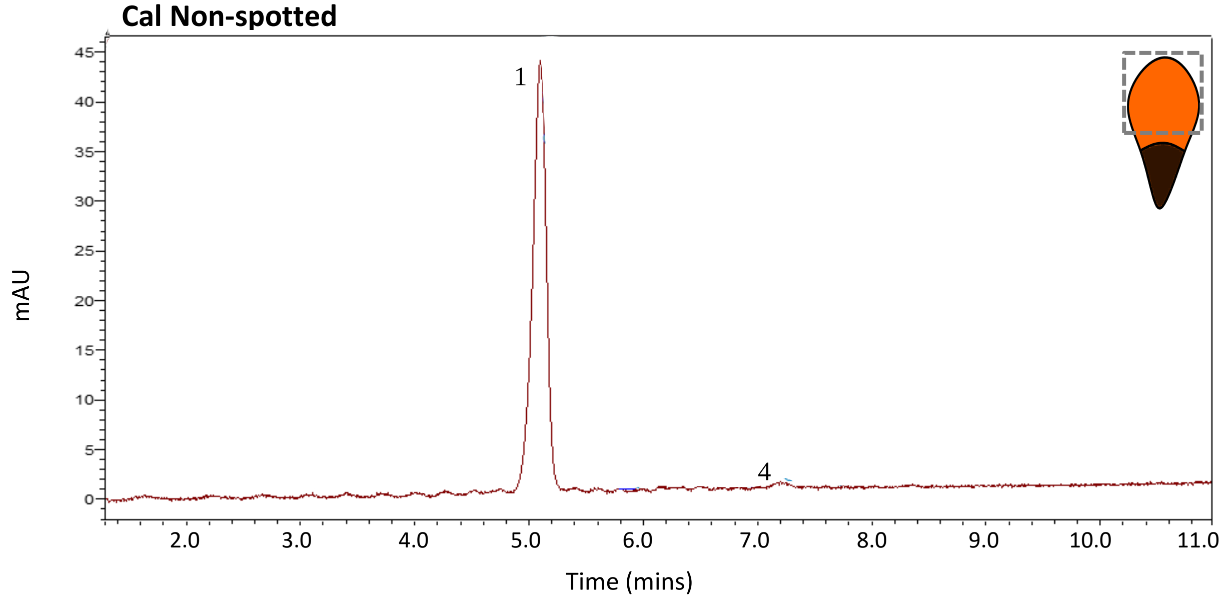
Cal Non-spotted
mAU
Time (mins)
1
4
0
5
10
15
20
25
30
35
40
45
2.0
3.0
4.0
5.0
6.0
7.0
8.0
9.0
10.0
11.0

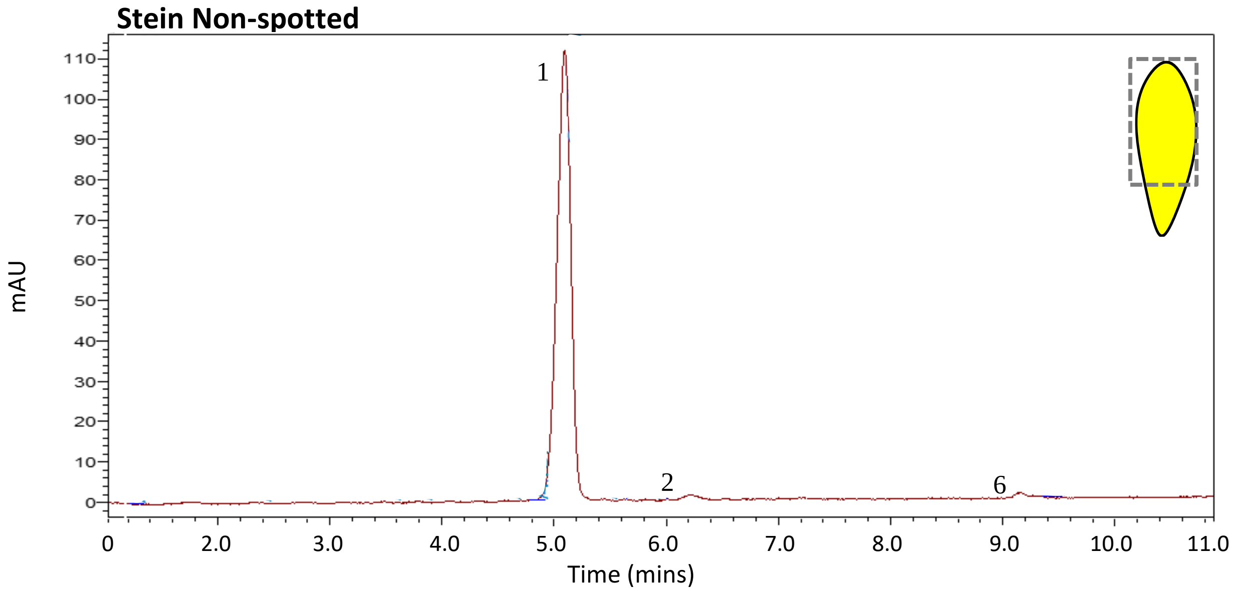
Stein Non-spotted
mAU
Time (mins)
1
2
6
0
10
20
30
40
50
60
70
80
90
100
110
0
2.0
3.0
4.0
5.0
6.0
7.0
8.0
9.0
10.0
11.0

Ultra-high-performance liquid chromatography (UHPLC-MS/MS) chromatograms of *G. diffusa* ray floret tissue at 525nm (bandwidth 50nm) – within the absorbance spectra of anthocyanins. Corresponding peaks are numbered throughout, with the MS spectra of each listed in Table S3. The tissue sample collected for each analysis is indicated by the grey box on the ray floret diagrams.

**Fig. S4** Mass spectra from *G. diffusa* UHPLC-MS/MS

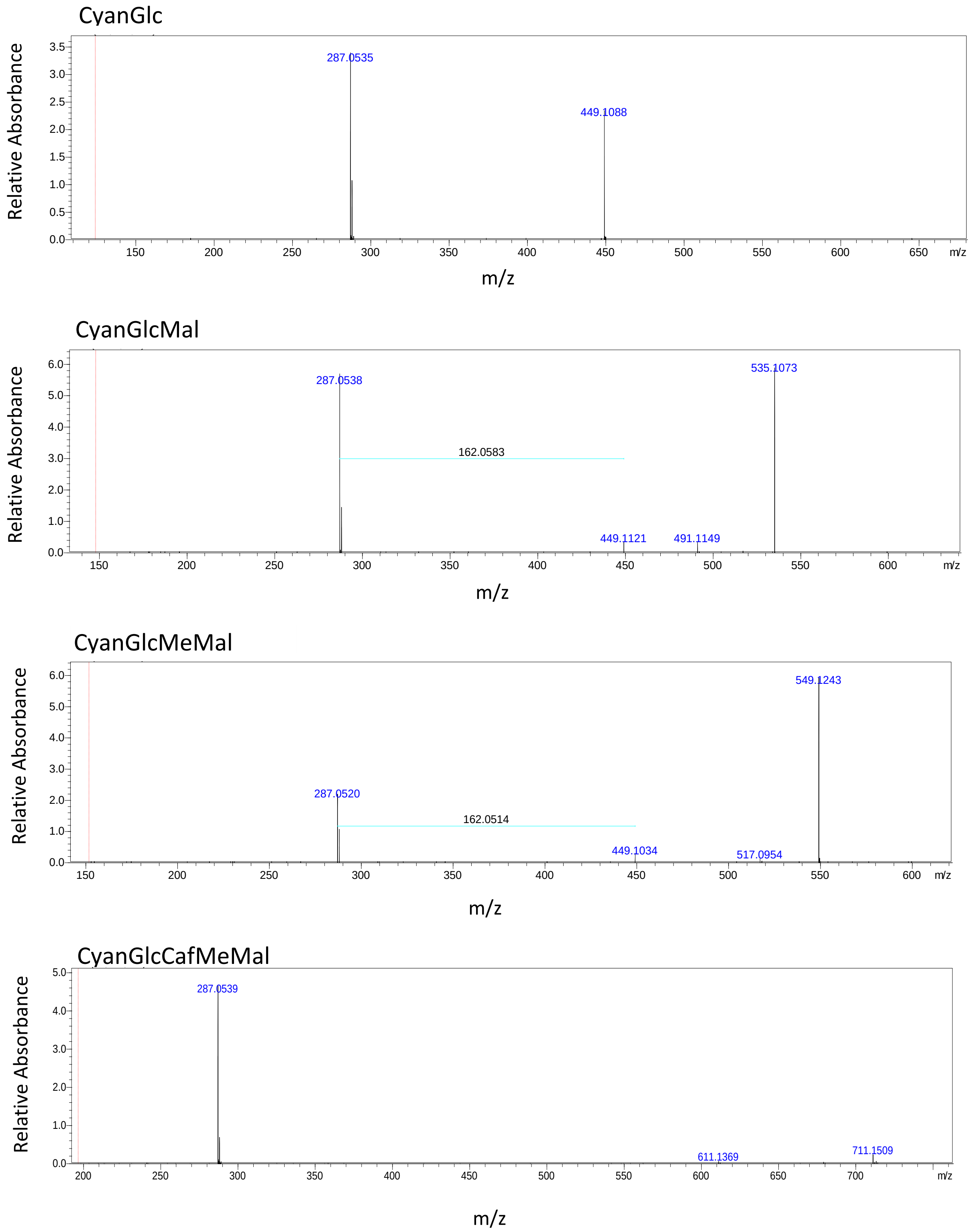


Example of mass spectra (MS2) used to identify anthocyanins. The mass spectra shown are from several peaks identified from the *G. diffusa* ‘Spring spot’ samples. Cyan = cyanidin, Glc = glucoside, Caf = cafeate residue, Mal = malonyl residue, MeMal = methylmalonyl residue.

**Fig. S5** Ray floret developmental stages used in qRT-PCRs in the morphotypes Spring, Cal, and Stein

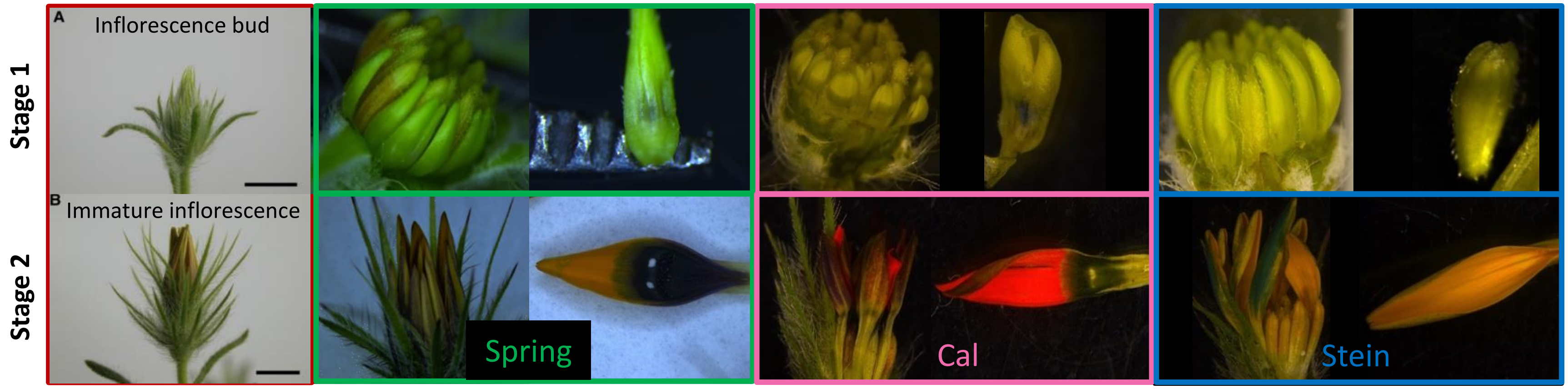


To assess whether there was equivalence in the timing of spot development between morphotypes, Spring and Cal were examined at two different developmental time points. Developmental stage 1 (top row) was defined as the time during which ray floret height was within the range of being equal to – x 1.5 the height of developing disc florets. At developmental stage 2 (bottom row) ray floret height was at least double that of developing disc florets, but ray florets were not yet fully mature.

At stage 1 the ray floret is yellow/green and the spot is a very small dark patch, by stage 2 the production of pigment on the abaxial side of the ray floret is initiating and the specialised cell types of the spot are initiating. Each Cal ray floret was cut into two latitudinally to give a non-spotted top segment and a spotted bottom segment. In Spring, the spot occupies a highly variable proportion of the ray floret and at developmental stage 1 the initiating spot is enclosed by the ray floret peripheral petals. As such, the Spring tissue used was whole non-spotted ray florets and whole spotted ray florets. All Spring individuals with a noticeable 'mark' (simple spot at the base of the non-spotted ray floret petals) were excluded to prevent this from confounding the 'spotted' versus 'non-spotted' comparison. Whole ray florets from Stein plants were used for expression analyses and only Stein individuals that did not produce spots were used. The red panels indicate the outward appearance of the buds at each developmental stage. The green, pink and blue panel provide close up views of the ray floret and petals at both developmental stages for Spring, Cal and Stein, respectively.

**Fig. S6** Gene expression analysis in the non-spotted petals of the Stein morphotype

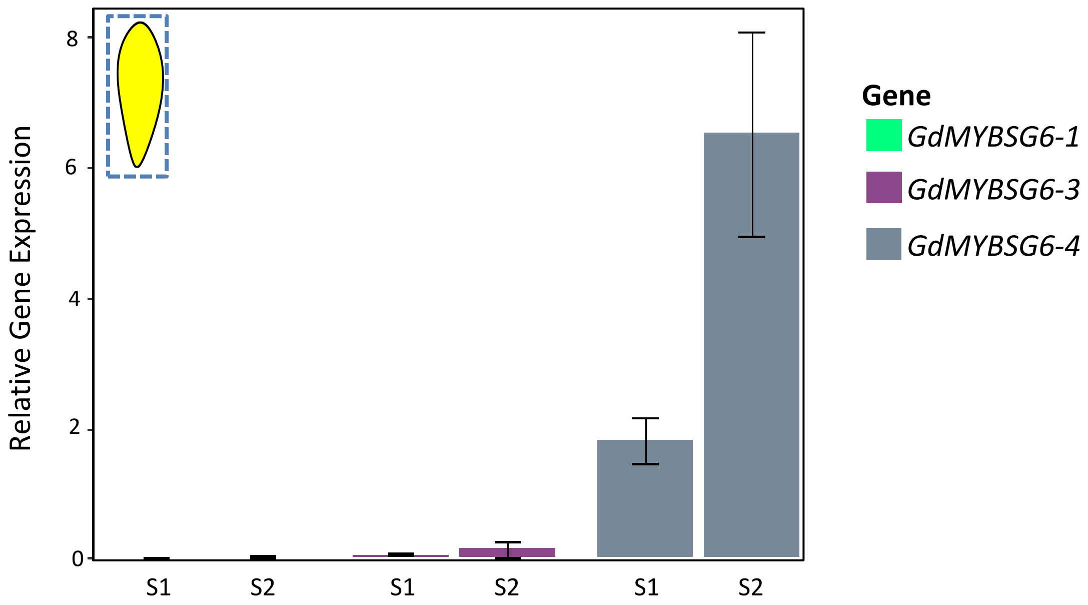


qRT-PCR results showing the relative expression of each *GdMYBSG6* gene at two developmental stages (S1 and S2) in Stein. *GdMYBSG6-2* is yet to be fully characterised in Stein so its expression was not investigated here. The entire petal tissue was used, as indicated in the ray floret diagrams. Error bars represent mean ± s.e.

**Fig. S7** Gene expression analysis in the Cal and Spring morphotypes

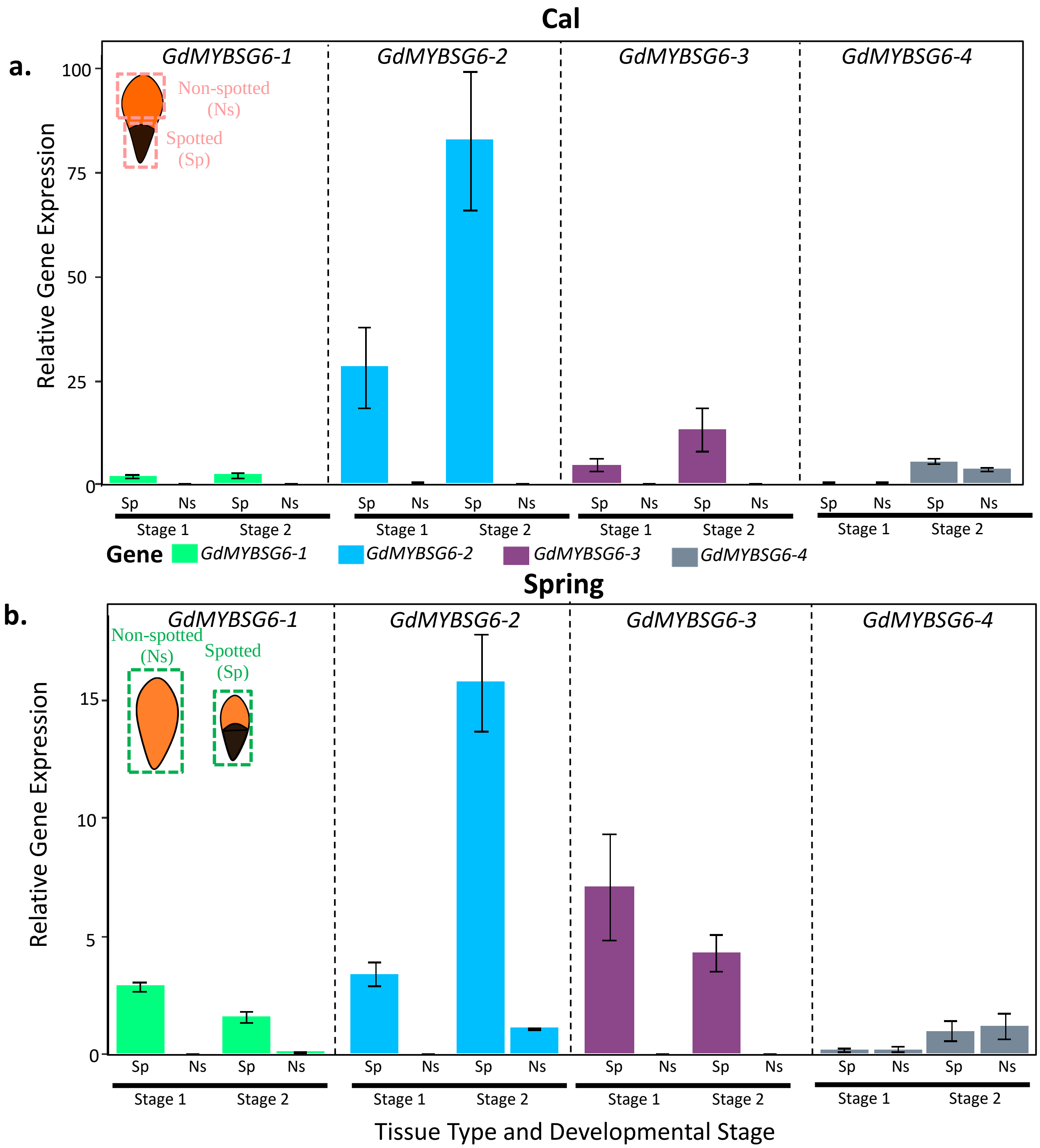


qRT-PCR results showing the relative expression of each *GdMYBSG6-1*, *2*, *3* and *4* gene at two developmental stages in a) Cal spotted (Sp) and non-spotted (Ns) ray floret petal tissue. b) Spring whole spotted ray florets (Sp) and whole non-spotted ray florets (Ns). The tissue segments used are indicated in the ray floret diagrams for each morphotype. Error bars represent mean ± s. e. The sample size for each tissue type was n = 3. For *GdMYBSG6-1*, *2*, and *3* spotted tissue had significantly higher expression levels than non-spotted tissue at both developmental stages. Only biologically relevant comparisons of expression levels were analysed through statistical tests.

**Fig. S8** Verification of transgene expression in transformed *N. tabacum* using RT-PCR

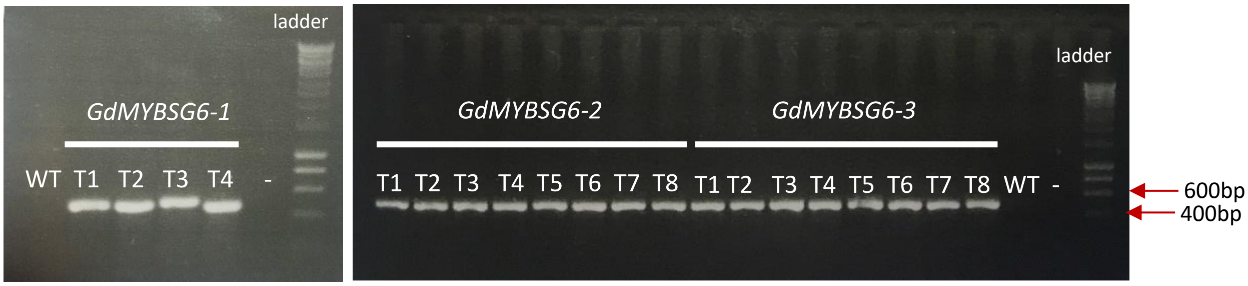


Gel electrophoresis images of PCR amplification of cDNA demonstrating that the relevant *GdMYBSG6* transgene is being expressed in each transgenic tobacco line in the $T_1$ generation. WT is wild type *N. tabacum* and the negative control for the PCR is indicated (-). Primers used to amplify *GdMYBSG6-2* and *GdMYBSG6-3* transgenes are the same, hence there is a single negative control and wild type sample present. Expected band sizes were as follows: *GdMYBSG6-1* 478bp, *GdMYBSG6-2* 520bp, and *GdMYBSG6-3* 549bp.

**Fig. S9** Developmental stages used for qRT-PCR and pigment extraction in *N. tabacum*

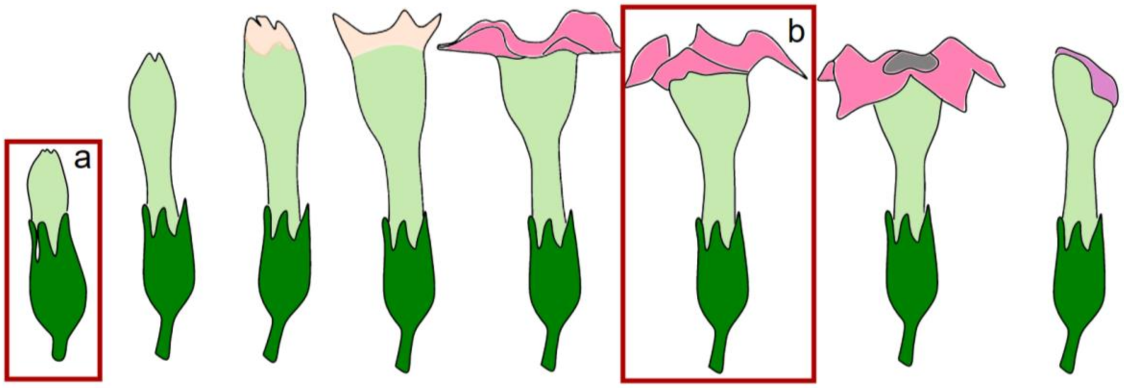


The developmental stages of wild type *N. tabacum* flowers (Pak Dek et al. 2017). The red boxes indicate the stages used in the analyses: (a) for qRT-PCR samples, (b) for anthocyanin extraction samples.

**Fig. S10** Leaves of *N. tabacum* transgenic lines constitutively overexpressing *GdMYBSG6* genes

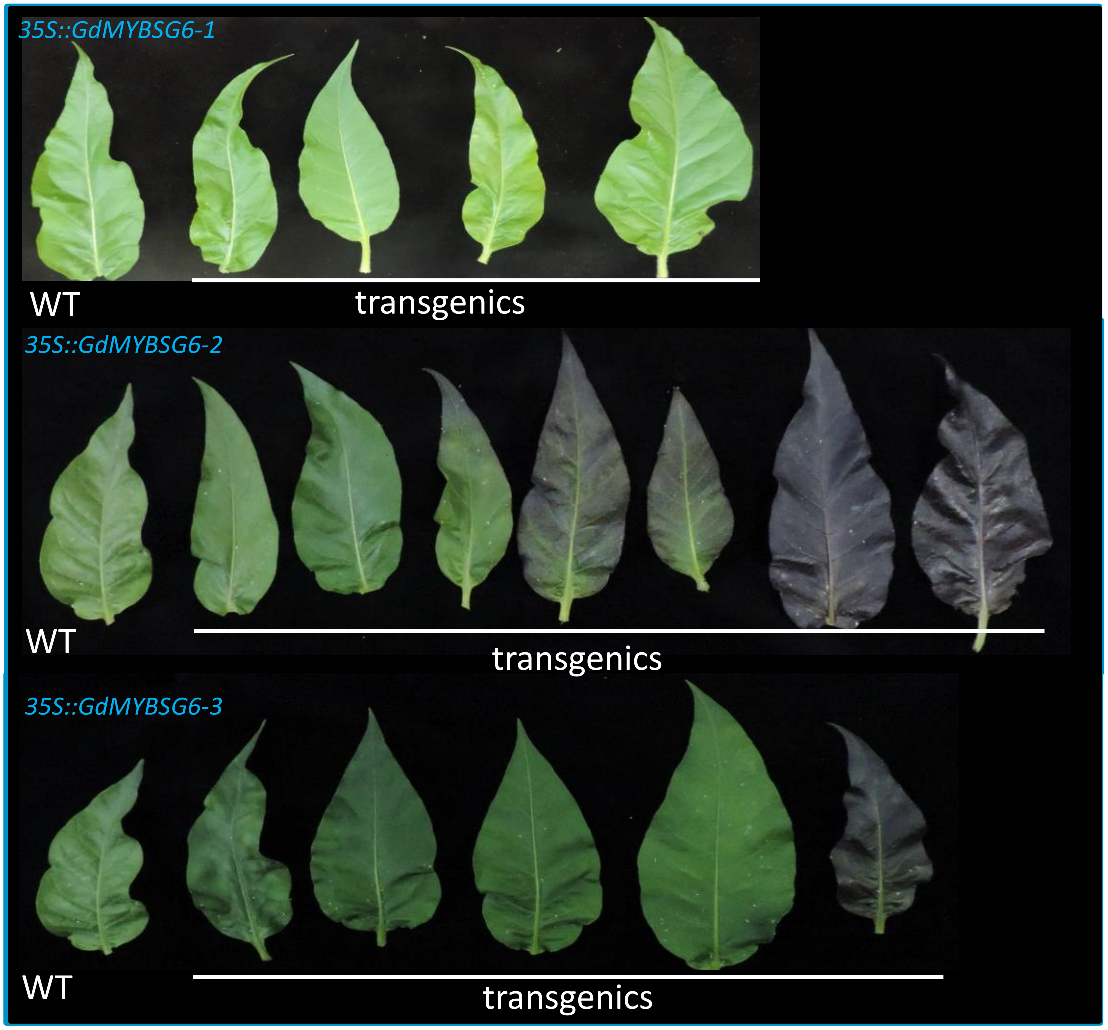


Leaves from T1 *Nicotiana tabacum* plants stably transformed with either *GdMYBSG6-1, GdMYBSG6-2*, or *GdMYBSG6-3* under the control of a double *35S* promoter and a wild type (WT) plant. Each leaf is taken from a $T_1$ individual, each originating from an independent $T_0$ line. Some anthocyanin pigmentation was eventually visible in the *35S::GdMYBSG6-1* lines compared to WT during leaf senescence (data not shown).

**Fig. S11** Anthocyanin extractions from floral tissue of transgenic *N. tabacum* lines constitutively overexpressing *GdMYBSG6* genes

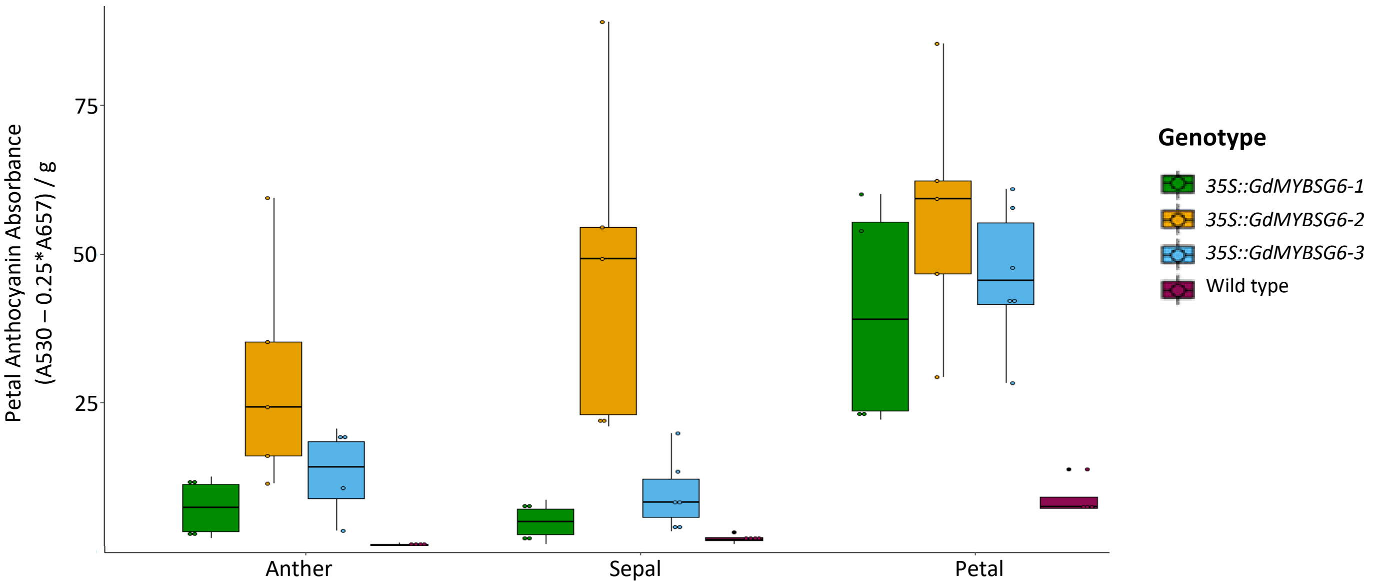


Anthocyanin content in $T_1$ *Nicotiana tabacum* plants stably transformed with either *GdMYBSG6-1*, *GdMYBSG6-2*, or *GdMYBSG6-3* under control of a strong constitutive double 35S promoter. The anthocyanin absorbance within anthers, petals, and sepals is given. Each data point is represented by a black circle and is the mean anthocyanin content in one independent line calculated from 3 - 6 samples.

**Fig. S12** Chromatograms of anthocyanin petal extractions from transgenic *N. tabacum* constitutively overexpressing *GdMYBSG6* genes

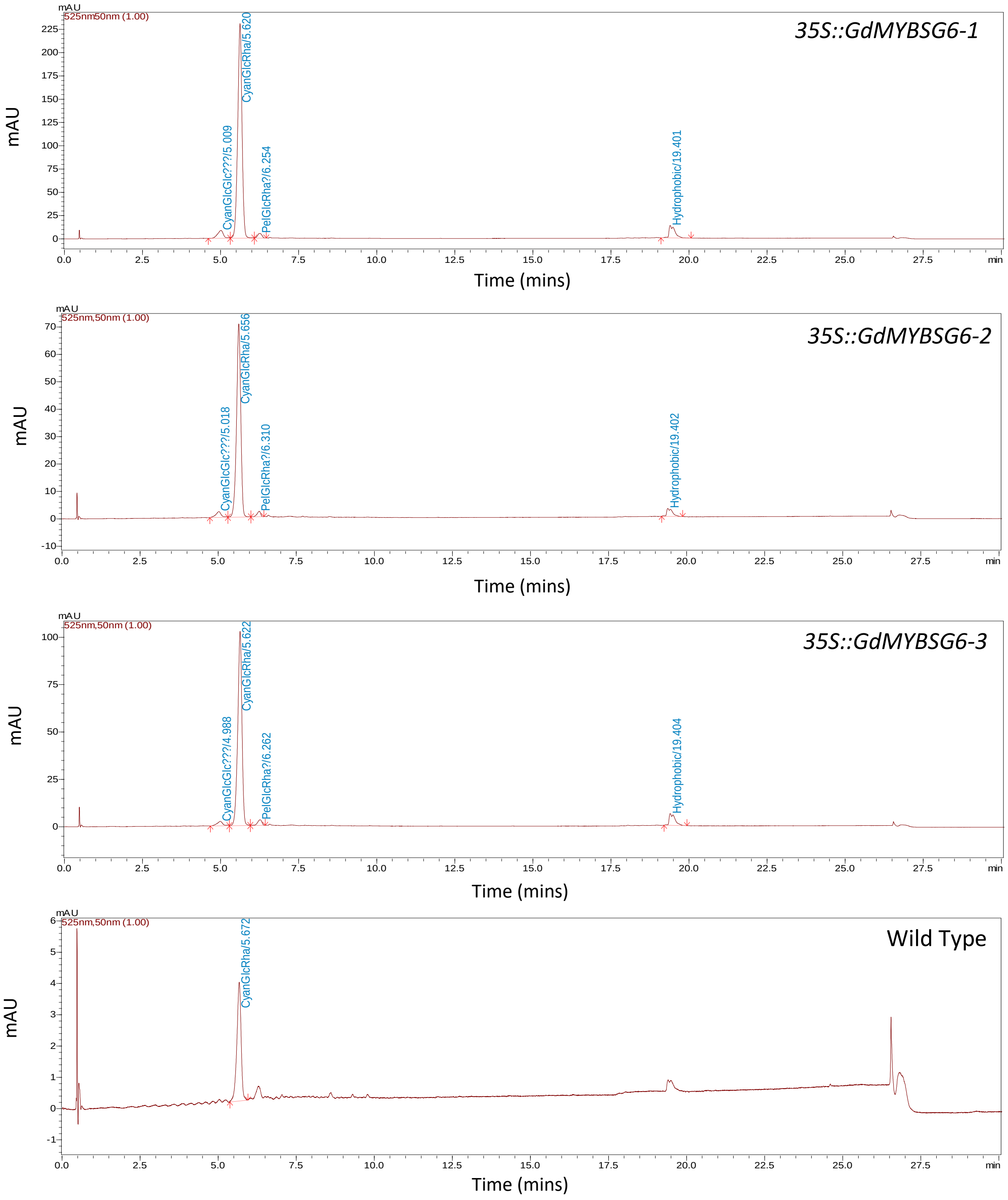


Ultra high performance liquid chromatography (UHPLC-MS/MS) chromatograms of *N. tabacum* petal tissue from a wild type plant and plants transformed with either *35S::GdMYBSG6-1*, *35S::GdMYBSG6-2*, or *35S::GdMYBSG6-3* at 525nm (bandwidth 50nm) – within the absorbance

spectra of anthocyanins. Corresponding peaks were identified using the MS spectra shown in Figure S13.

**Fig. S13** Spectra of anthocyanin petal extractions from transgenic *N. tabacum* constitutively overexpressing *GdMYBSG6* genes

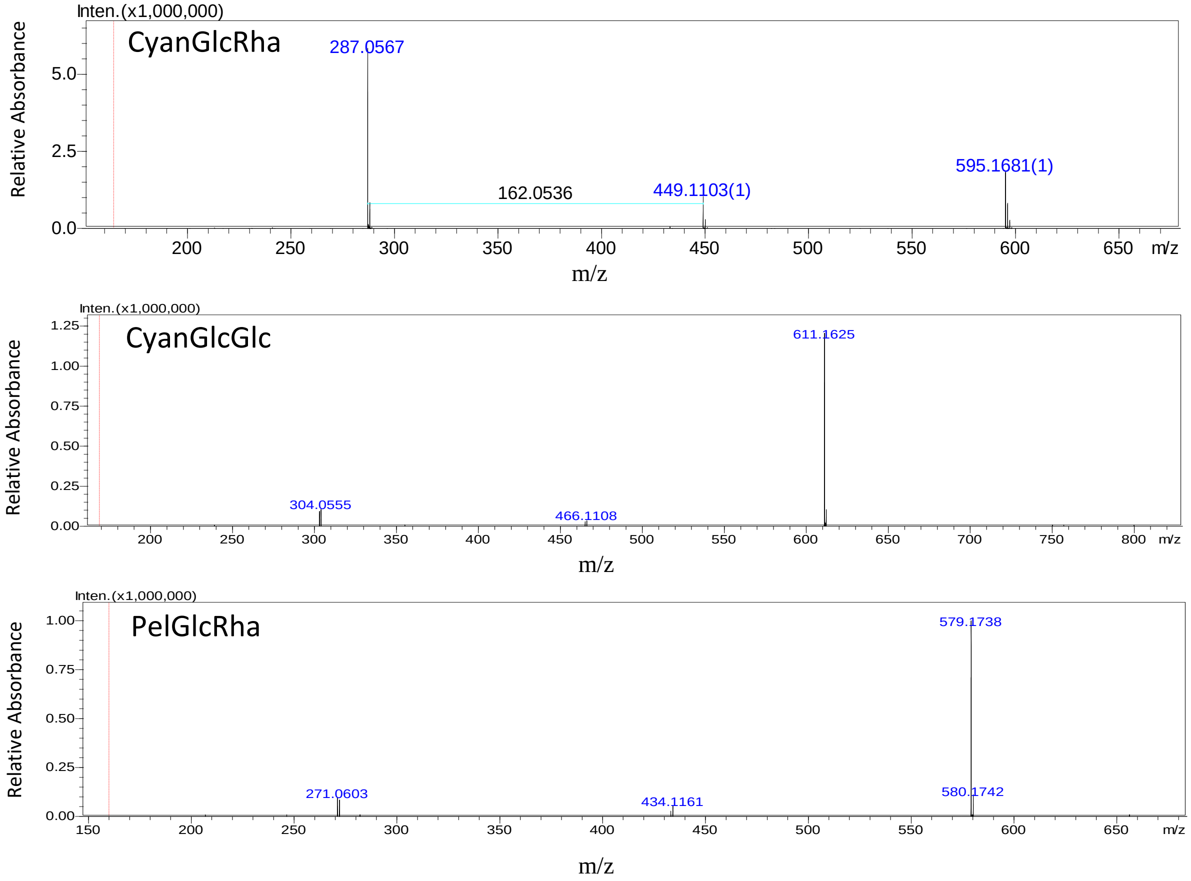


Mass spectra (MS2) used to identify anthocyanins. The mass spectra shown are from peaks identified in a *N. tabacum* sample. Cyan = cyanidin, Glc = glucoside, Pel = pelargonidin, Rha = rhamnose.

**Fig. S14** *GdMYBSG6* relative expression levels in floral tissue of transgenic *N. tabacum* carrying a *35S::GdMYBSG6* transgene

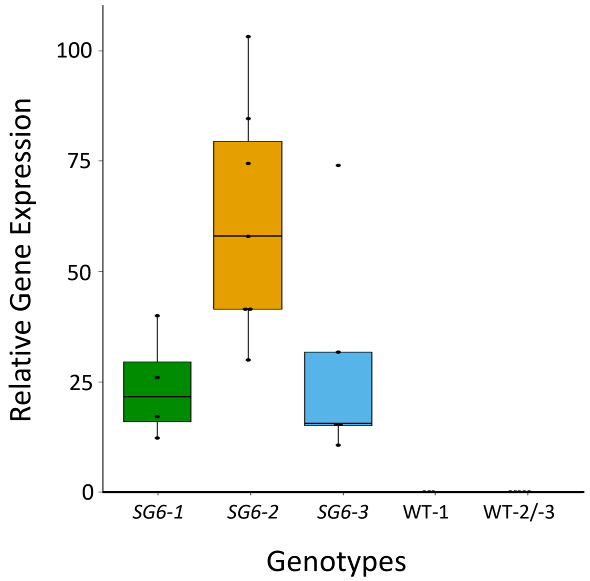


Relative expression of the transgene within several lines for each set of transformants. Primers used to assess transgene expression levels were the same for *GdMYBSG6-2* and *GdMYBSG6-3*. Wild type expression levels are shown for each set of primers (WT-1 with *GdMYBSG6-1* primers and WT-2/-3 with *GdMYBSG6-2/-3* primers). The black line in each box indicates the median value and the whiskers 25/75% quantile +/- 1.5 * interquartile range, respectively. Each data point (black dot) is from a $T_1$ individual, each originating from an independent $T_0$ line.

**Fig. S15** Expression analysis of *G. diffusa* genes coding for late anthocyanin synthesis enzymes

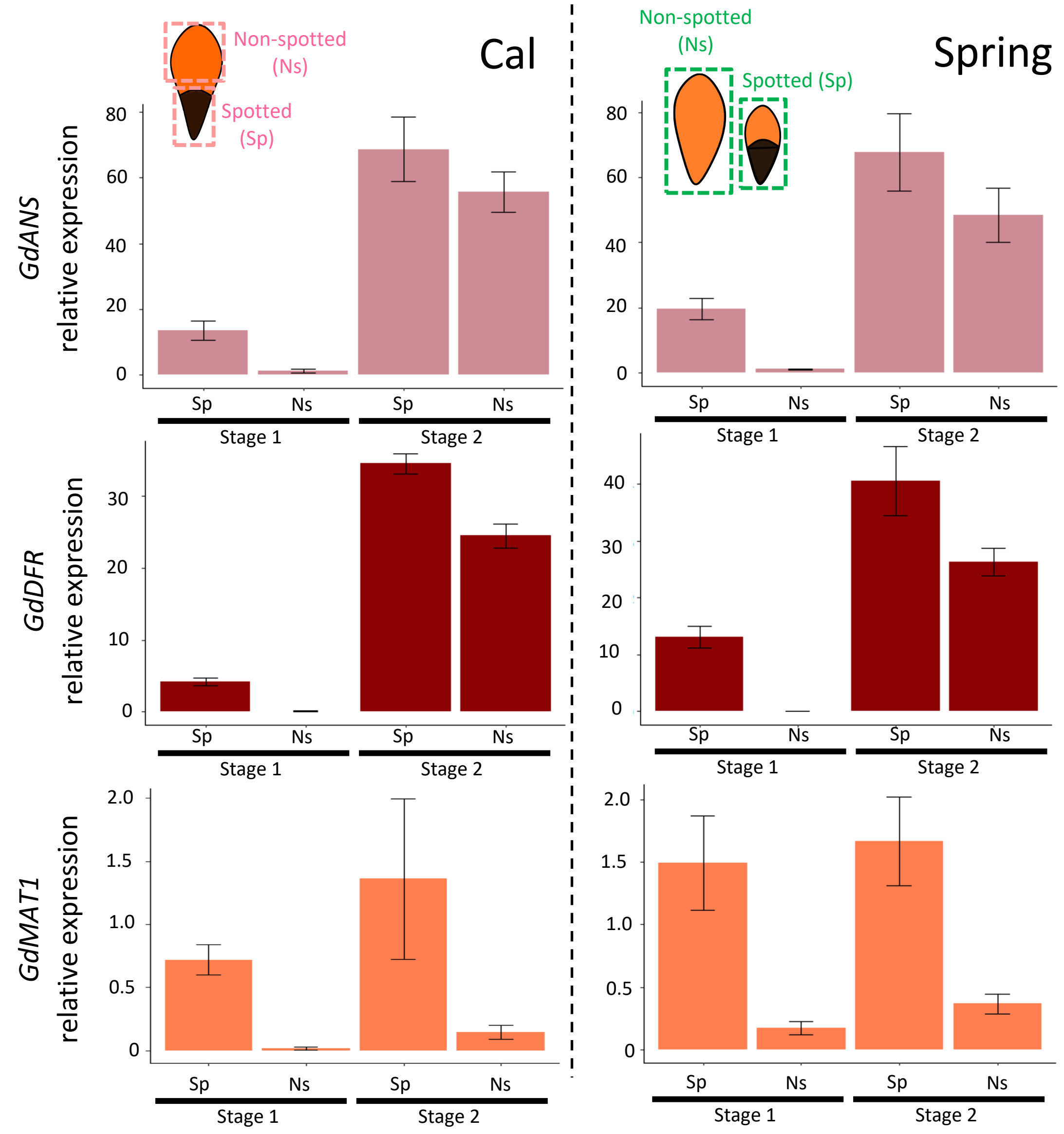


qRT-PCR results showing the relative expression levels of genes encoding the anthocyanin synthesis enzymes: anthocyanidin synthase (ANS), dihydroflavonol 4-reductase (DFR) and malonyl transferase (MAT1) in *G. diffusa* morphotypes Spring and Cal. Gene expression is relative to the reference gene *Elongation Factor 2* (*GdEF-2*). Two developmental stages were used in Cal and Spring morphotypes: Cal spotted (Sp) and non-spotted (Ns) ray floret petal tissue (left-hand column) and Spring whole spotted ray floret petals (Sp) and whole non-spotted ray floret petals (Ns) (right-hand column). These were sampled at two developmental stages (1) and (2). The tissue segments used are indicated in the ray floret diagrams for each morphotype. Error bars represent the mean ± s. e of three biological replicates.

**Table S1** Primer sequences

| Name | Amplified product description | Sequence |
|---|---|---|
| **GdActin.F** | Reference gene for checking cDNA synthesis has worked | CCAAGGGCAGTGTTTCCTAGT |
| **GdActin.R** | Reference gene for checking cDNA synthesis has worked | TGGTACGACCACTGGCATAG |
| **GeneRacer.R** | 3'RACE to amplify 3'UTRs | GCTGTCAACGATACGCTACGTAACG |
| **GeneRacer.Rn** | 3'RACE nested primer to amplify 3'UTRs | CGCTACGTAACGGCATGACAG |
| **M13.F** | Used in colony PCR to amplify insert in the pBLUE vector | GTAAAACGACGGCCAGT |
| **M13.R** | Used in colony PCR to amplify insert in the pBLUE vector | CAGGAAACAGCTATGAC |
| **GdEF-2.qP.F** | Reference gene for qRT-PCR in Spring | CAACTGCAGCGGGTCCATTAT |
| **GdEF-2.qP.R** | Reference gene for qRT-PCR in Spring, Cal, and Stein | CAGCTGTCATAACTTGCCCTGAA |
| **GdEF-2C.qP.F** | Reference gene for qRT-PCR in Cal and Stein | ACTGCAGCGGGTCCATTATGT |
| **GdSG6-1.qP.F** | *GdMYBSG6-1* for qRT-PCR in Spring and Stein | TCTACTCAAAACACCAAATGATGATCTT |
| **GdSG6-1.qP.R** | *GdMYBSG6-1* for qRT-PCR in Spring, Cal, and Stein | AACTTCGGTGCCAGTGTT |
| **GdSG6-1C.qP.F** | *GdMYBSG6-1*for qRT-PCR in Cal | CCTACTCAAAACACCAAATGATGATCTT |
| **GdSG6-2.qP.F** | *GdMYBSG6-2* for qRT-PCR in Spring and Cal | AAACACGAAAAGCAAACGGACA |
| **GdSG6-2.qP.R** | *GdMYBSG6-2* for qRT-PCR in Spring | AAAAGATAGGGTTAGGAACATCAACGT |
| **GdSG6-2C.qP.R** | *GdMYBSG6-2* for qRT-PCR in Cal | GGTTAGGAACATCAACGGTGTGAA |
| **GdSG6-3.qP.F** | *GdMYBSG6-3* for qRT-PCR in Spring, Cal, and Stein | TTCTTTGGTGGAGAGGCAGG |
| **GdSG6-3.qP.R** | *GdMYBSG6-3* for qRT-PCR in Spring, Cal, and Stein | GATAGGGGTAGGAACATCAAAGTTGTT |
| **GdSG6-4.qP.F** | *GdMYBSG6-4* for qRT-PCR in Spring | ACAAAATGTCCCCAACTTTAATCTCGTC |
| **GdSG6-4.qP.R** | *GdMYBSG6-4* for qRT-PCR in Spring | GACCACCCAATTTCACAGTCAAATTCATC |
| **GdSG6-4C.qP.F** | *GdMYBSG6-4* for qRT-PCR in Cal and Stein | GTGGTCATTTGGTGGTTCTTCGA |
| **GdSG6-4C.qP.R** | *GdMYBSG6-4* for qRT-PCR in Cal and Stein | TTCCGAATGTAGCAAGTCCCACATG |
| **GdANS.qP.F** | *GdANS* for qRT-PCR in Spring and Cal | CGTTCCCGGAGGAGAAAC |
| **GdANS.qP.R** | *GdANS* for qRT-PCR in Spring and Cal | GTGGCGAGTGCTCGTAG |
| **GdDFR.qP.F** | *GdDFR* for qRT-PCR in Spring and Cal | GCGAAAACAGTCAAGAGGCTAGTT |
| **GdDFR.qP.R** | *GdDFR* for qRT-PCR in Spring and Cal | AATGTCCCTCATCGTAAACAGGAAGT |
| **GdMAT1.qP.F** | *GdMAT1* for qRT-PCR in Spring, Cal, and Stein | CCATCTTTTCTTCTACGAATTCCCCTACTC |
| **GdMAT1.qP.R** | *GdMAT1* for qRT-PCR in Spring, Cal, and Stein | TGACACCTGAATTATTAGGGTTGGA |

| Name | Amplified product description | Sequence |
|---|---|---|
| **GdSG6-1.F** | Used to amplify full length *GdMYBSG6-1* | GAAATAGAAATGAGCATGTACTTC |
| **GdSG6-1.R** | Used to amplify full length *GdMYBSG6-1* | CAATCTCAATCTTTATGCAAC |
| **GdSG6-2.F** | Used to amplify full length *GdMYBSG6-2* | GAATTTCATCTTTGTCTTCTACA |
| **GdSG6-2.R** | Used to amplify full length *GdMYBSG6-2* | CAACTTCCATTCATCTTGGA |
| **GdSG6-3.F** | Used to amplify full length *GdMYBSG6-3* | CAATAAAAACGAGCATGTACATC |
| **GdSG6-3.R** | Used to amplify full length *GdMYBSG6-3* | AATATGAGTAAAAGATAGGGGTAGGA |
| **GdSG6-4.F** | Used to amplify full length *GdMYBSG6-4* | ACATGATAGGCTCCTCCCATCTG |
| **GdSG6-4.R** | Used to amplify full length *GdMYBSG6-4* | CTAGAGACAATTATTTCACAGGAATCTGAG |
| **NtEF1.qP.F** | *NtEF-1* reference gene for qRT-PCR in *N. tabacum* | TGAGATGCACCACGAAGCTC |
| **NtEF1.qP.R** | *NtEF-1* reference gene for qRT-PCR in *N. tabacum* | CGTTAAACCCAACATTGTCACCAG |
| **NtUBC.qP.F** | *NtUBC* reference gene for qRT-PCR in *N. tabacum* | GCAGCACGCATGTTCAGTGA |
| **NtUBC.qP.R** | *NtUBC* reference gene for qRT-PCR in *N. tabacum* | CAGTCTGCTGTCCAGCTCTG |
| **GdSG6-1.tob.qP.F** | *GdMYBSG6-1* amplifying transgene for qRT-PCR in *N. tabacum* | AGGACAAGACACCACAGTCACA |
| **GdSG6-1.tob.qP.R** | *GdMYBSG6-1* amplifying transgene for qRT-PCR in *N. tabacum* | CCTGAACCATTATGAACCCATTTAGGT |
| **GdSG6-23.tob.qP.F** | *GdMYBSG6-2* and *GdMYBSG6-3* amplifying transgene for qRT-PCR in *N. tabacum* | CACAAGACACCACGGTCACA |
| **GdSG6-23.tob.qP.R** | *GdMYBSG6-2* and *GdMYBSG6-3* amplifying transgene for qRT-PCR in *N. tabacum* | TGACCCATCATGAACCCATTTAGGT |
| **NtANS.qP.F** | *NtANS* gene for qRT-PCR in *N. tabacum* | TCCTCCACAATATGGTGCCTG |
| **NtANS.qP.R** | *NtANS* gene for qRT-PCR in *N. tabacum* | GGGTGTCCCCAATATGCATGA |
| **NtDFR.qP.F** | *NtDFR* gene for qRT-PCR in *N. tabacum* | ACTGAGTTTAAAGGCATCGATAAGGACT |
| **NtDFR.qP.R** | *NtDFR* gene for qRT-PCR in *N. tabacum* | TGAATTGAAACCCCATATCCGTCAG |
| **GdANS.F** | Used to amplify full length *GdANS* | CACAACAAAACCACAAACAC |
| **GdANS.R** | Used to amplify full length *GdANS* | CAAAGAGCAACACTAATGTGATG |
| **GdMAT1.F** | Used to amplify full length *GdMAT1* | CACCATCCTCTCTCAACCAATTCA |
| **GdMAT1.R** | Used to amplify full length *GdMAT1* | CTCGAAACAAATCAAAACCAATCA |
| **GdDFR1.F** | In 5' UTR of *GdDFR* to amplify into gene | ACACTCACCACTCACCAGT |
| **GdDFR2.F** | At start codon of *GdDFR* to amplify into gene | AAATGAAAGAGGATTCTCCTACCAC |

| Name | Amplified product description | Sequence |
|---|---|---|
| **GdDFR1.R** | In 3'UTR of one *GdDFR* 'variant' to amplify into gene | CTTGATTTTATTGACTTGAACCA |
| **GdDFR2.R** | In 3'UTR of several *GdDFR* 'variant' to amplify into gene | TACAAACCCCTGCCACATC |
| **GdDFR3.R** | In 3'UTR of one *GdDFR* 'variant' to amplify into gene | CACCGTTTGTAATCTTTTCATTTACAG |
| **GdDFR4.R** | In 3'UTR of one *GdDFR* 'variant' to amplify into gene | GAGCACCATTTGTAATCTTATTATGTAGA |
| **GdDFR5.R** | In 3'UTR of one *GdDFR* 'variant' to amplify into gene | TGACCATCAACGTTTTTGACAGAA |
| **GdDFR6.R** | In 3'UTR of all *GdDFR* 'variant' to amplify into gene | GCACCATTTGTAATCTTGTCATTTAGAG |
| **Nt.geno.S1.F** | Used to check transgene expression in *GdMYBSG6-1* transgenic *N. tabacum* | AGGACAAGACACCACAGTCACA |
| **Nt.geno.S23.F** | Used to check transgene expression in *GdMYBSG6-2* and *GdMYBSG6-3* transgenic *N. tabacum* | CACAAGACACCACGGTCACA |
| **Nt.geno.R** | Used to check transgene expression in transgenic *N. tabacum,* in transcribed portion of 35S terminator | TTATCGGGAAACTACTCACACA |
| **PR.GdDFR.F** | For amplifying *GdDFR* promoter region | AGAAACCATGTTACTTGTTACGACA |
| **PR.GdMAT1.F** | For amplifying *GdMAT1* promoter region | GTAACCGCTTTCTACCTTCTATCTCTCTT |
| **PR.GdANS.F** | For amplifying *GdANS* promoter region | TCCGTGTCAAAAATGGATATAAAAAGTAC |
| **GP.SG6-4.F** | Degenerate primer to amplify *G. personata MYBSG6-4* | ATGAGACMNGGTAGTAATAAGGNN |
| **GP.SG6-4.R** | Degenerate primer to amplify *G. personata MYBSG6-4* | TCAAAGTTGTTCCGAATGTAGCNN |
| **GP.SG6-4.F** | Internal primer to amplify *G. personata MYBSG6-4* | GGACCGCCGAAGAAGACAA |
| **GP.SG6-4.R** | Internal primer to amplify *G. personata MYBSG6-4* | CCCCGTCCATAGGGAAATCAA |
| **GdDFR.deg1.F** | Degenerate primer to isolate the promoter region of *GdDFR* | CCAYYGTNTGYGTCACH |
| **GdDFR.deg2.F** | Degenerate primer to isolate the promoter region of *GdDFR* | MGTYATGAGACTNCTYSAAC |

**Table S2** Anthocyanin quantification and characterisation

| Compound | Spring Spot | Cal Spot | Spring Mark | Spring Top | Spring Non-spotted | Cal Non-spotted | Stein Non-spotted |
|---|---|---|---|---|---|---|---|
| **CyanGlc** | 0.3610 ± 0.0132 | 0.1331 ± 0.0139 | 0.1087 ± 0.0084 | 0.0244 ± 0.0011 | 0.1796 ± 0.0009 | 0.4551 ± 0.0083 | 0.3452 ± 0.0007 |
| **CyanGlcMal isomer** | 0.0156 ± 0.0008 | 0.0048 ± 0.0005 | 0.0038 ± 0.0004 | not detected | 0.0010 ± 0.0005 | 0.0010 ± 0.0004 | 0.0071 ± 0.0007 |
| **CyanPentose** | 0.0033 ± 0.0003 | trace | 0.0005 ± 0.0002 | trace | trace | 0.0014 ± 0.0007 | 0.0007 ± 0.0001 |
| **CyanGlcMal** | 0.5051 ± 0.0121 | 0.1676 ± 0.0121 | 0.1164 ± 0.0063 | 0.0014 ± 0.0006 | 0.0040 ± 0.0007 | 0.0196 ± 0.0068 | trace |
| **CyanGlcMeMal** | 0.1346 ± 0.0186 | 0.0693 ± 0.0021 | 0.0422 ± 0.0029 | 0.0005 ± 0.0004 | trace | 0.0035 ± 0.0019 | not detected |
| **CyanGlcCaf** | 0.0040 ± 0.0008 | trace | not detected | not detected | not detected | trace | 0.0021 ± 0.0004 |
| **CyanGlcCafMal** | 0.0266 ± 0.0039 | trace | 0.0028 ± 0.0006 | trace | not detected | trace | trace |
| **CyanGlcCafMeMal** | 0.0043 ± 0.0018 | not detected | 0.0013 ± 0.0002 | not detected | not detected | not detected | trace |
| **Total anthocyanins** | 1.0544 ± 0.4550 | 0.3761 ± 0.0945 | 0.2755 ± 0.0409 | 0.03* | 0.1852 ± 0.0727 | 0.4810 ± 0.3445 | 0.3559 ± 0.1981 |

Types of anthocyanin detected in *G. diffusa* ray floret tissue through UHPLC-MS/MS. Cyan = cyanidin, Glc = glucoside, Caf = cafeate residue, Mal = malonyl residue, MeMal = methylmalonyl residue, Pentose = pentose sugar. Each value represents the approximate anthocyanin concentration (µg/mg) of each compound (± s.e., n = 3), calculated by multiplying the proportion of anthocyanin the compound represents with the total anthocyanin content. Trace is used where only 1/3 samples contained the compound and the mean relative quantity was <0.0006. *Estimated from total peak areas relative to other samples and their total anthocyanin contents. 'Spring Top' is the top non-spotted section of the Spring spotted ray floret.

**Table S3** Anthocyanins detected in the ray floret tissue of *G. diffusa*

| Compound | Retention time (min) | Identity | m/z | HPLC-ESI(+) - MS experiment m/z |
|---|---|---|---|---|
| 1 | 5.059 | CyanGlc | 449 | MS2 [449]: 287* |
| 2 | 6.281 | CyanGlcMal isomer | 535 | MS2 [535*]: 287, 401 |
| 3 | 6.484 | CyanPentose | 535 | MS [535]: 240, 287, 331, 403*, 419, 426, 449, 466 |
| 4 | 7.133 | CyanGlcMal | 535 | MS2 [535*]: 287, 449, 491 |
| 5 | 8.283 | CyanGlcMeMal | 549 | MS2 [549*]: 287, 449, 517 |
| 6 | 9.109 | CyanGlcCaf | 611 | MS2 [611]: 231, 258, 287*, 333, 373, 487, 606 |
| 7 | 10.827 | CyanGlcCafMal | 697 | MS2 [697]: 287*, 493, 585 |
| 8 | 12.183 | CyanGlCafMeMal | 711 | MS2 [711]: 287*, 611 |

UV spectra and physical properties of all anthocyanins found in the ray floret tissue of *G. diffusa* through UHPLC and positive mode electrospray mass spectrometry. Compound number corresponds to peak number in 1.15, * indicates the base peak. Cyan = cyanidin, Glc = glucoside, Caf = caffeate residue, Mal = malonyl residue, MeMal = methymalonyl residue, Pentose = pentose sugar.

**Methods S1** Anthocyanin quantification

Walker (2012) found that the ray florets of Spring and Cal contain chlorophyll. Consequently, the absorbance measurements of each pigment extraction were taken at both A530 (the peak of anthocyanin absorption) and A657 (the peak of absorption of chlorophyll degradation products in acidic methanol). Dilution factor was accounted for and the total anthocyanin content calculated using the equation $A = A530 - 0.25 * A657$. (Mancinelli 2020). Total anthocyanin content (A) was divided by the fresh weight of the sample to give (B), which was used to calculate approximate absolute anthocyanin concentrations of sample extracts (µg/mg fresh weight): concentration=(B/34) x 484.83 (Airoldi et al. 2019), where 484.83 is the molecular weight of cyanidin 3-glucoside and 34 is used as the millimolar extinction coefficient (Gerats et al. 1982).

| time (minutes) | % acetonitrile |
|---|---|
| 0.01 | 2 |
| 0.50 | 2 |
| 5.00 | 10 |
| 17.00 | 30 |
| 25.00 | 90 |
| 25.80 | 90 |
| 26.00 | 2 |
| 30.10 | 2 |

Gradient of acetonitrile versus 1% (v/v) formic acid in water used during UHPLC-MS/MS analysis.

**Methods S2** Map of plasmids used in *N. tabacum* transformation

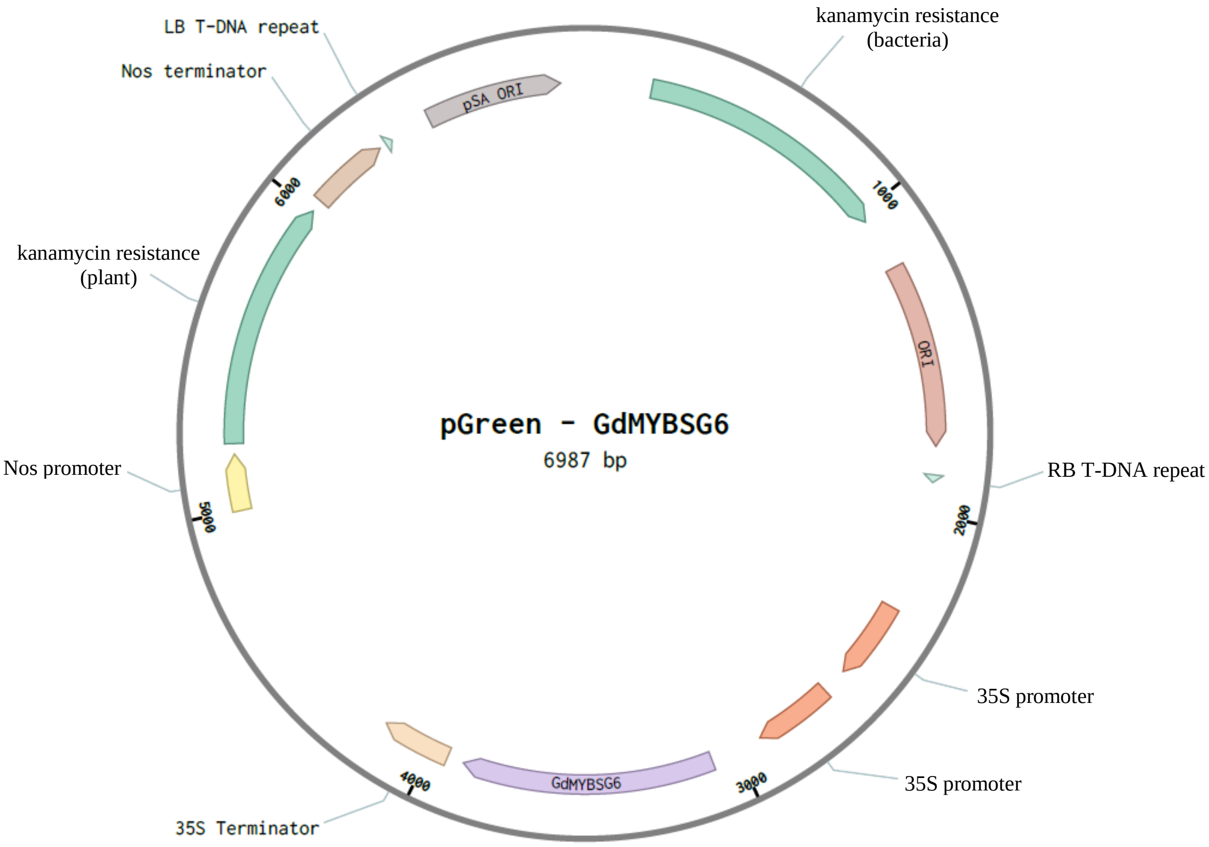


Map of the pGreen-*35S::GdMYBSG6* plasmids used to stably transform *N. tabacum*. Three plasmids were used differing only in the *GdMYBSG6* coding sequence that they contained, either *GdMYBSG6-1*, *GdMYBSG6-2*, or *GdMYBSG6-3*.

**Methods S3** *N. tabacum* stable transformation procedure

Young *N. tabacum* leaves were soaked in 10% bleach solution, rinsed in DI water and submerged in a suspension of transformed *A. tumefaciens*. Leaves were cut into segments while submerged, transferred to filter paper to remove excess *A. tumefaciens* and placed onto an MS9 plate (0.5mg/ml IAA and 1mg/ml BAP). These plates were incubated for 48 hrs in the dark at room temperature. Leaf discs were then transferred to new MS plates (0.5mg/ml IAA, 1mg/ml BAP, 200µg/ml ampicillin, 100µg/ml kanamycin, and 500µg/ml cefotaxime). Every ten days plant tissue was transferred onto fresh MS plates, containing the hormones and antibiotics listed above. Once 1 - 2cm shoots had sprouted from the calli that leaf disks produced, these were cut at the base and transferred to Hamilton jars containing MS9 (200µg/ml ampicillin, 100µg/ml kanamycin, and 500µg/ml cefotaxime). Once a sufficient root network had grown the plants were transferred to the greenhouse and potted in Levington's M3 bedding compost.

The following media was used during transformation and regeneration of *N. tabacum*:
***Agrobacterium* suspension**: 2.2 g/l Murashige-Skoog Medium with vitamins (Duchefa), 35 g/l sucrose, ddH2O
**MS9 media**: 4.4 g/l Murashige-Skoog Medium with vitamins (Duchefa), 20 g/l sucrose, ddH2O
**MS media**: 4.4 g/l Murashige-Skoog Medium with vitamins (Duchefa), 30 g/l sucrose, ddH2O

**Hormone and antibiotic concentrations:**
MS plates leaf discs initially transferred to: 0.5 mg/ml IAA and 1 mg/ml BAP.
MS plates leaf discs transferred to after 48 hrs: 0.5 mg/ml IAA, 1 mg/ml BAP, 200 mg/l ampicillin, 100 mg/l kanamycin, and 500 μg/ml cefotaxime.
Hamilton jars containing MS9 media shoots were transferred to: 200 mg/l ampicillin, 100 mg/l kanamycin, and 500 mg/l cefotaxime.

**Methods S4** Map of plasmid used to produce the recombinant GdMYBSG6-2 protein used in EMSA

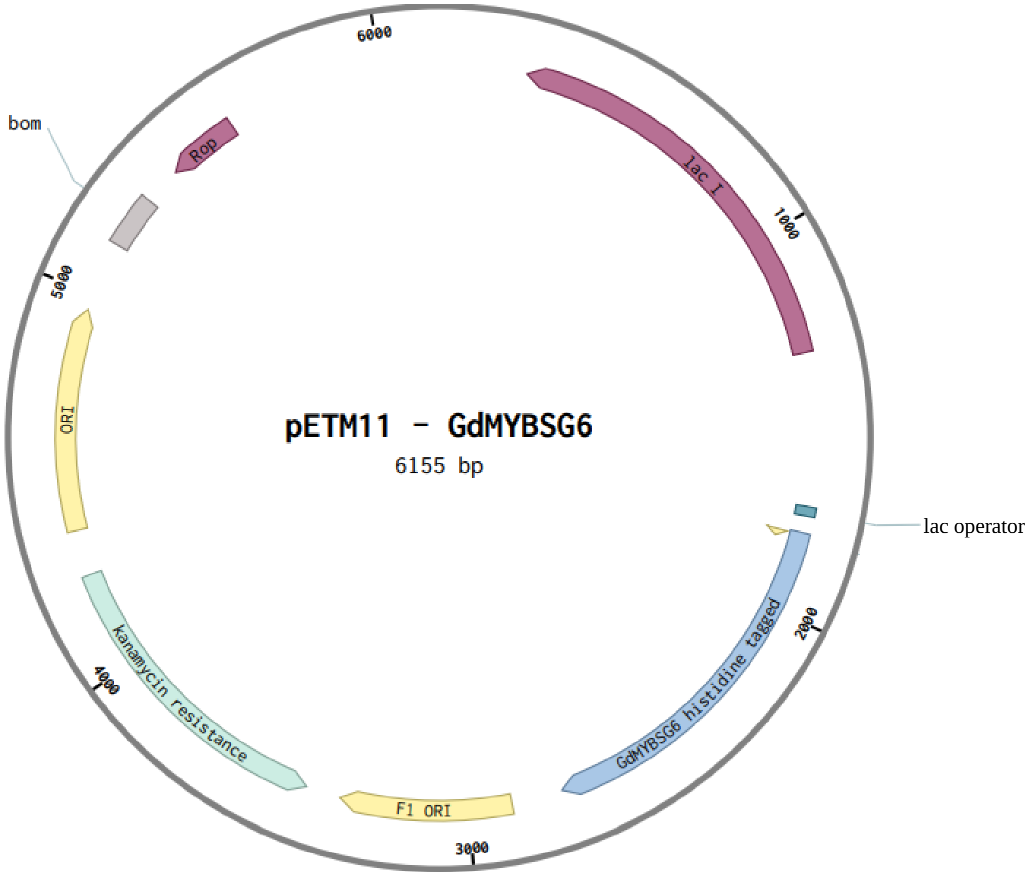


Map of the pETMII–*GdMYBSG6-2* plasmid introduced into the rosetta II strain of *E. coli*. The *GdMYBSG6-2* coding sequence was fused in N-term and C-term to histidine encoding nucleotides.

Protein production was induced in the rosetta II strain of *E. coli* and histidine tagged protein produced.

**Methods S5** Buffer solutions used for recombinant protein purification and EMSA

Base buffer: 5ml Tris 1M pH8 (final 10mM), 25ml NaCl 5M (final 250mM), 25ml glycerol (final 5% v/v), $ddH_2O$ up to 0.5L.

Lysis buffer: 24ml base buffer, 50µl DTT 1M (final 2mM), 0.25g sarkosyl (final 1%), 250µl PMSF 100mM (final 1mM).

Loading buffer: 48ml base buffer, 100µl DTT 1M (final 2mM), 0.5g sarkosyl (final 1%), 17.4mg imidazole (final 5mM).

Filtration buffer 1: 49ml base buffer, 100µl DTT 1M (final 2mM), 0.25g sarkosyl (final 0.5%), 109mg imidazole (final 40mM).

Filtration buffer 2: 25ml base buffer, 50µl DTT 1M (final 2mM), 0.125g sarkosyl (final 0.5%), 425mg imidazole (final 250mM).

Acrylamide gel destaining solution: 50ml glacial acetic acid, 150ml ethanol, 300ml $ddH_2O$.

Protein storage buffer (dialysis buffer): 100µl Tris 1M pH8 (final 10mM), 500µl NaCl 5M (final 250mM), 500µl glycerol (final 5% v/v), 20µl DTT 1M (final 2mM), $ddH_2O$ up to 10ml.

10x annealing buffer to generate dsDNA used in EMSA: 100mM tris pH7.5, 1.5M NaCl, 10mM EDTA pH8.

Binding buffer (pH 7.5) for protein-DNA reactions used in EMSA: 1ml 1M tris pH8, 1.5ml 5M NaCl, 25µl 0.5M EDTA pH8, 100µl 1M
MgCl2, igepal (NP-40) 10µl, 10% (v/v) 1M DTT, ddH2O up to 50ml.

**Methods S6** Map of plasmid used in dual luciferase assays

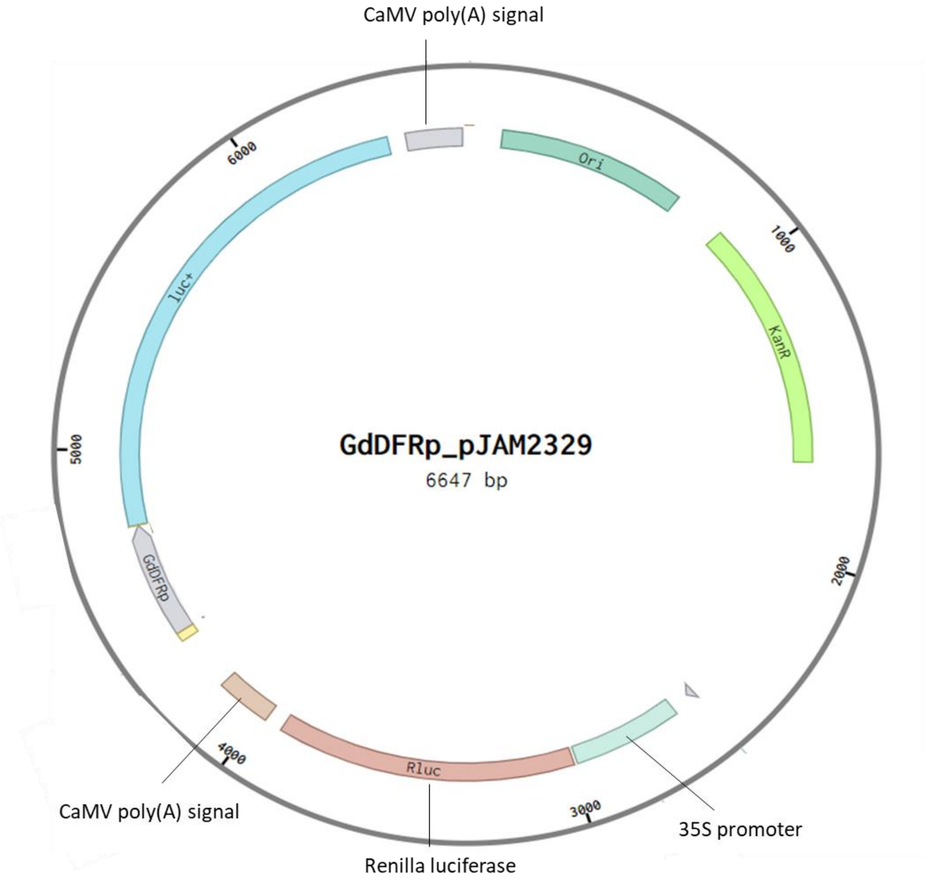


Map of the pJAM2329 – *GdDFRp* reporter construct used in dual luciferase assays. The *GdDFR* promoter region was fused to a luciferase reporter gene.
